# Deep learning reveals the complex genetic architecture of a highly polymorphic sexual trait

**DOI:** 10.1101/2023.09.29.560175

**Authors:** Wouter van der Bijl, Jacelyn J. Shu, Versara S. Goberdhan, Linley M. Sherin, Changfu Jia, María Cortázar-Chinarro, Alberto Corral-López, Judith E. Mank

## Abstract

The extraordinary variation in male guppy coloration has proven a powerful model for studying the interplay of natural and sexual selection. However, this variation has hampered the high-resolution characterization and determination of the genetic architecture underlying male guppy color, as well as clouded our understanding of how this exceptional level of diversity is maintained. Here we identify the heritability and genetic basis of male color variation using convolutional neural networks for high-resolution phenotyping coupled with selection experiments, controlled pedigrees and whole-genome resequencing for a Genome Wide Association Study (GWAS) of color. Our phenotypic and genomic results converge to show that color patterning in guppies is a combination of many heritable features, each with a largely independent genetic architecture spanning the entire genome. Autosomally-inherited ornaments are polygenic, with significant contributions from loci involved in neural crest cell migration. Unusually, our GWAS results suggest that gene duplicates from the autosomes to the Y chromosome are responsible for much of the sex-linked variation in color in guppies, providing a potential mechanism for the maintenance of variation of this classic model trait.

## Introduction

The study of color polymorphisms has been pivotal to our understanding of how variation is maintained in populations. Perhaps none are as polymorphic as the color patterns of male guppies, *Poecilia reticulata*, which have been extensively studied in the context of fitness^1–4^ and female preference^5,6^. The widespread variation in the extent and patterning of male color across wild populations^4,7,8^ and the rapid response to natural and artificial selection^9,10^, coupled with high heritability estimates^11,12^ all suggest that substantial genetic variation for this trait is maintained in guppy populations. These colors are not only unusually variable for sexual ornaments but have some of the highest coefficients of genetic variation ever reported for morphological traits^11,13^. Although female mate preference for rare male patterns likely acts to preserve some variation from the eroding forces of genetic drift and selection^5,13,14^, the genetic architecture underlying extraordinary diversity in male guppy color remains mysterious, as does the maintenance of this exceptional diversity.

Part of the difficulty in determining the genetic basis of male guppy coloration lies in the multiple forms of variation. Yellow-orange xanthophores and black melanophores form spots that vary among males in their location, as well as in the extent and saturation of these colors. Color pattern variation arises from embryonic neural crest cells that form the precursors of xanthophores and melanophores^15^. As the administration of testosterone can induce color traits in females, coloration appears to be under hormonal control^8,16^. Therefore, the genetic basis of color variation is expected to include loci involved in the migration of neural crest cells, as well as genes involved in the synthesis and deposition of pigments, and possibly the production of and response to testosterone. Importantly, orange carotenoids are not synthesized *de novo* in vertebrates, but are collected from food sources^17^ and modified prior to deposition, and so the genetic basis of orange coloration could also include loci involved in the bioavailability of these molecules.

It has often been suggested that, despite the striking diversity of guppy color patterns, their genetic basis might be surprisingly simple. However, the excessive complexity of guppy pattern phenotypes has significantly complicated the identification of the underlying genetic loci. Early studies, which assessed pattern inheritance by visual inspection, found a mix of Y-linked, X-linked, and autosomal inheritance for specific color traits^8,12,18–21^. More recently, a study suggested that the complete color patterns in one population are inherited as a single Y-linked locus with twenty-seven alleles^5^. These findings indicate that at least some ornaments are monogenic and co-inherited due to linkage, but they leave unresolved how ornaments with more complex inheritance patterns are controlled. More quantitative and objective summary pattern measures, such as the total orange or black area, suggest a mix of inheritance patterns^11,12,16,22^. Here we overcome these longstanding obstacles by developing a pipeline of convolutional neural networks for high-resolution phenotyping of 3,229 fish coupled with selection experiments and controlled pedigrees to objectively determine the heritability of the full complexity of color patterns. We then used our phenotype data for a Genome Wide Association Study (GWAS) of color to identify the genetic architecture of a suite of color traits.

Our phenotypic and genomic results converge to show that color patterning in guppies is a combination of many heritable features, each with largely independent genetic architectures. We delineate fifteen separate ornaments with heterogeneous genetic architectures, with some ornaments inherited as a single sex-linked locus, while others have a polygenic autosomal basis. Finally, our results suggest that gene duplications from the autosomes to the sex chromosomes are responsible for much of the sex-linked variation in color in guppies, providing a potential mechanism for the maintenance of variation of this classic model trait.

### Artificial selection on orange coloration

Starting from an outbred laboratory population with large variation in color patterning, we created three replicate pairs of selection lines for reduced and increased amounts of orange coloration as a percent of total body area. Each male in the experiment (*n* = 3,229) had multiple photos taken from each side to track measurement error (*n* = 14,100). Using a series of neural networks, we performed image segmentation to separate the fish from the background, and to identify the orange and black coloration on the fish (Fig 1A). In addition to orange, xanthophores also create the less frequent yellow color patches, which we include in our orange measurements. We then warped images to a common reference shape based on placed landmarks to enable fine-scale color mapping^12^. Combined with a full pedigree, this dataset allowed us to trace the inheritance of color patterns across four generations (Fig 1B). While some body areas are almost always pigmented (Fig 1C), color is rare in most places on the male body plan, giving each male a distinct pattern.

**Figure 1:**
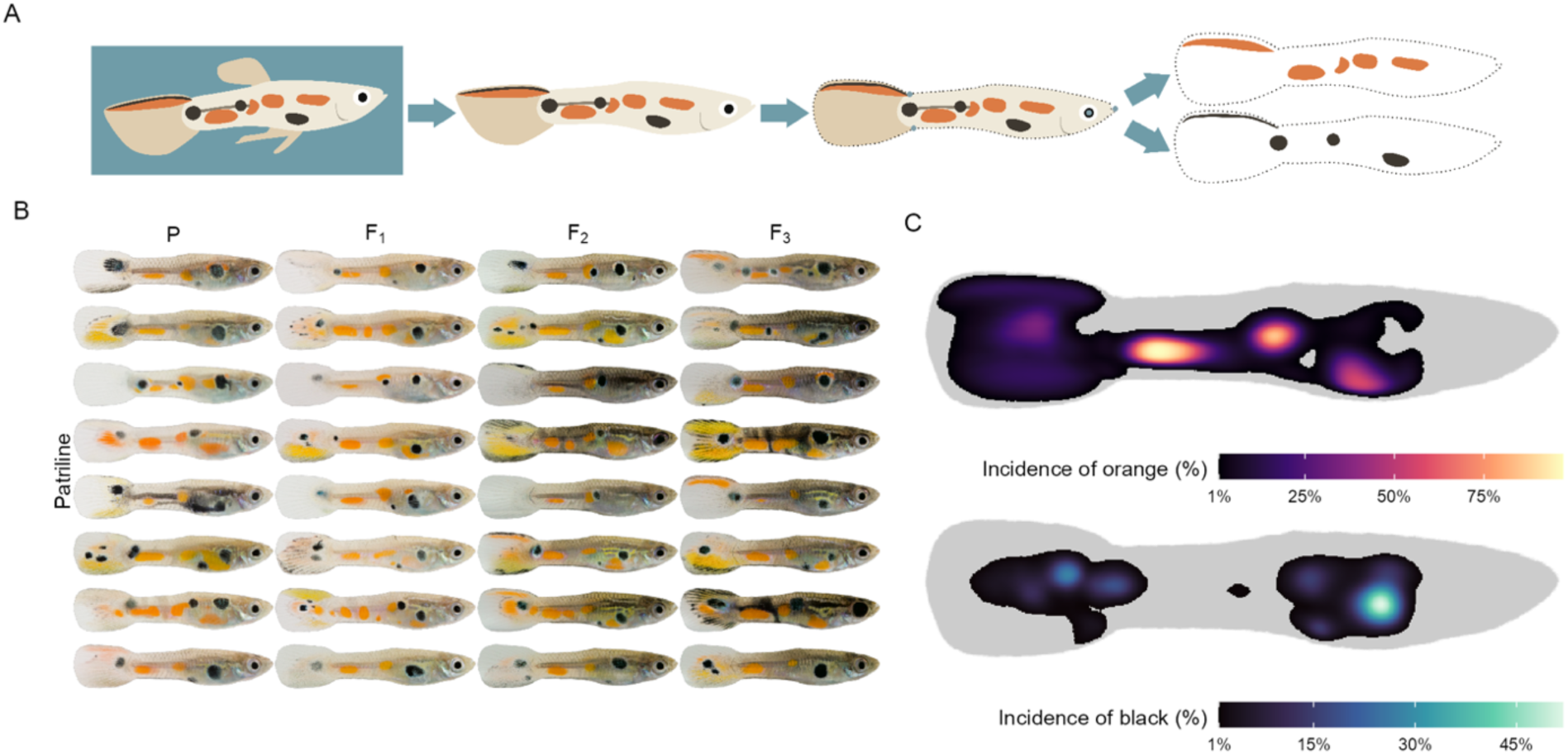
Quantification of pattern variation. A) Overview of the phenotyping pipeline. From each photo, the fish is extracted, the body is aligned to a common reference shape using landmarks, and the position of orange and black color is extracted. B) Randomly selected examples of color inheritance along patrilines. Images on the same row belong to the same patriline, and columns are successive generations of the selection experiment (both selection directions). C) The incidence of color across the guppy body among all males. Body positions where the incidence of color is less than 1% are colored grey.

The percent of orange color responded rapidly and strongly to selection (Fig 2A). After just three generations, up-selected males had more than double the amount of orange color compared to their down-selected counterparts (% up – % down in F_3_ [95% credible interval]: 6.05 [5.67, 6.48]). We modeled the inheritance of black and orange color along the pedigree^23^, separating autosomal, X-linked and Y-linked genetic effects, as well as influences due to the maternal and tank environments. The strong response in orange color to selection is consistent with its substantial heritability (narrow-sense *h*^2^: 0.59, [0.50, 0.68], Fig S1). While orange and black ornaments share the same body plan, selection on orange nonetheless had a negligible effect on the amount of black coloration (up – down in F_3_: -0.09 [- 0.17, 0.01], Fig S2A). Although the amount of black color is also substantially heritable (*h*^2^: 0.67 [0.53, 0.80], Fig S3), the amount of the two colors is inherited largely independently as evidenced by their low genetic correlations (*r*_autosomal_: 0.07 [-0.15, 0.26], Fig S4). Although previous work on other populations^11,24–26^ has identified strong Y-linkage, our estimates of Y effects are low (*h*^2^_Y-linked_ orange: 0.03 [0.00, 0.09], black: 0.04 [0.00, 0.08]), and instead the genetic variation for the proportion of orange and black color on the body has an autosomal and, to a lesser extent, X-linked origin (Figs S1&S2).

**Figure 2:**
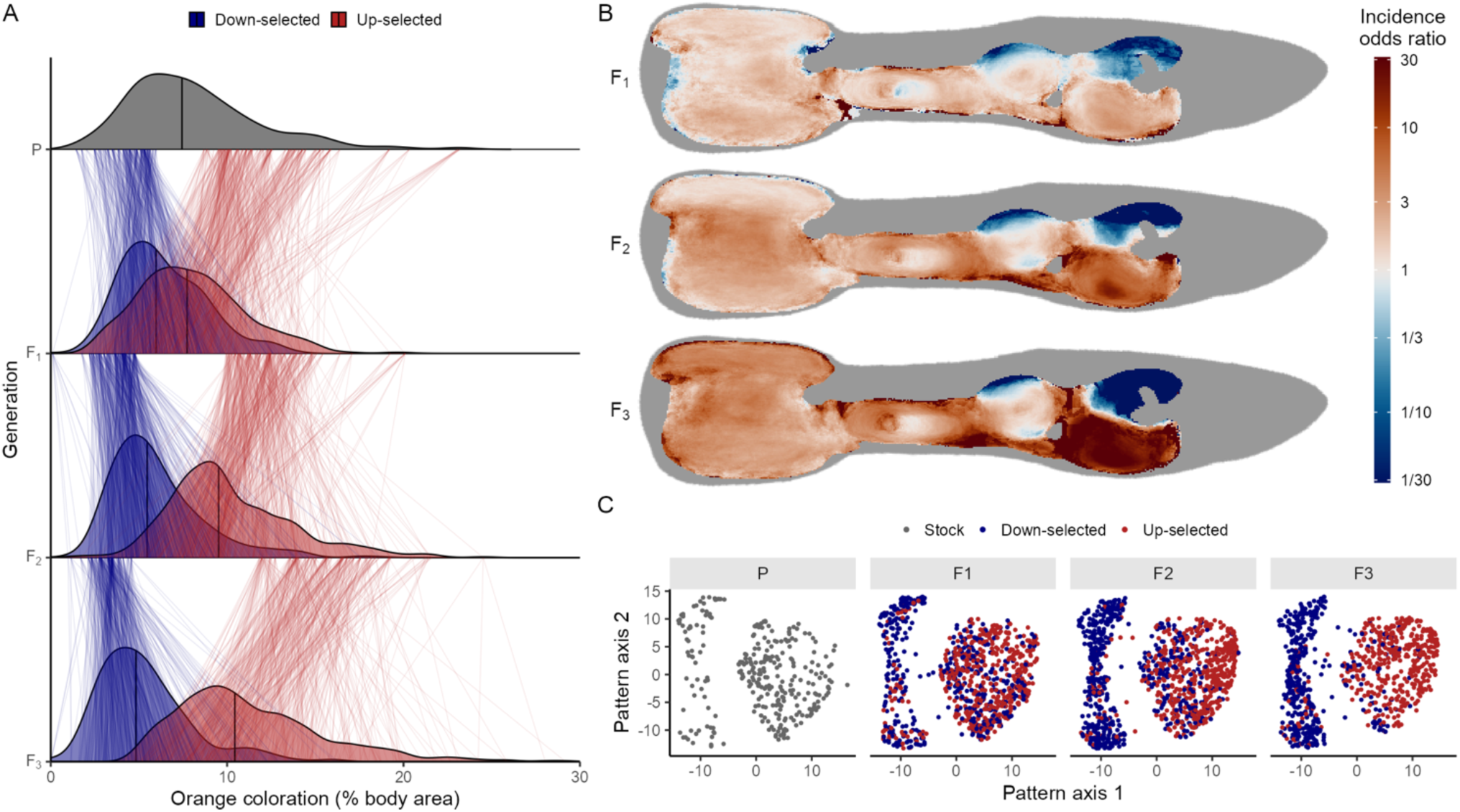
Orange color responds rapidly to selection. A) Density plots show the distribution of the percent of orange coloration per generation and selection regime. Thin lines between generations connect fathers to their sons. Thick lines inside the densities show the median value of color area. B) Heatmaps illustrating the effect of selection across the body. Each heatmap cell is colored by the log odds ratio (as estimated by a generalized linear mixed model), illustrating the relative odds that a male has orange color at that location. Body positions where the incidence of orange color is less than 1% are colored grey. C) Selection moves populations through orange patternspace. Axes show orange patternspace (after UMAP dimension reduction), with points indicating individuals, colored by selection direction, and facets showing the four generations.

Our selection regime affected the incidence of orange color at nearly all positions on the body, with effect sizes increasing with each successive generation of selection (Fig 2B). The strength and direction of the effects were spatially heterogeneous. In the F_3_ generation, up-selected males were approximately 5x more likely to have orange color at any position in the tail or caudal fin (posterior to the gonopodium). The strongest effects were located close to the pectoral fin and gills, where up-selected males were more than 100x as likely to have orange color on the ventral side, while surprisingly, they were more than 100x *less* likely to be orange dorsally (although this dorsal color was much less prevalent overall, Fig 1C). The response against the direction of selection at some locations, and correlated local changes in black color (Fig S2B), illustrate that some elements of the color pattern are genetically linked.

To investigate whether other traits were co-selected in our selection lines, we looked at changes in behavior and life history, since selection on embryonic neural crest cells has resulted in changes in these traits in other vertebrate systems^27,28^. Guppy males flaunt their coloration during characteristic sigmoid display behavior, where they curve their body into an S-shape, quiver, and swim laterally in front of females^29,30^. Using a subset of 126 male offspring from the F3 generation (63 from each selection direction, evenly split across the replicate lines), we evaluated the effects of our selection regime on male sexual behavior. In a standard no-choice test with a non-receptive female, males from the up-selected lines performed both more (short displays: ratio up/down = 1.74, z = 2.15, p = 0.031, long displays: up/down = 1.84, z = 2.28, p = 0.022) and longer (up/down = 1.9, z = 2.44, p = 0.015) displays to the female (Fig S5). In addition, they spent more time following the female (up/down = 1.9, z = 2.44, p = 0.015, Fig S6), and performed more attempts at coercive sneak copulations (up/down = 1.99, z = 2. 09, p = 0.037), suggesting the more colorful males were 1.7-2x more sexually active overall (Fig S6). Males did not show changes in body size or tail size (standard length in mm, up – down in F_3_: -0.36 [-0.84, 0.12], tail length: 0.10 [-0.21, 0.43], tail area mm^2^: 0.019 [-2.5, 2.2], Fig S7), nor did selection alter the fecundity of females (F_2_ offspring per brood, down-selected: 7.74, up-selected: 7.79, ratio up/down: 0.99 [0.87, 1.12]) or the time between pairing and the first brood (F_2_, up - down in days: -1.23 [-3.4, 0.97], Fig S8).

### Localized heritability, ornaments, and patternspace

While the quantity of orange and black coloration are important to female choice^7,31^, male patterning is largely inherited as discrete spots and stripes at fixed body positions^12,24–26^. To understand the spatial structure of color inheritance, we modeled the heritability of the presence of color separately for each body location (Fig 3), confirming the fine-scale heritability of color patterning. In many places the localized heritability is high (> 0.8), much higher than the heritability of the total amount of color. Decomposing the genetic variance into autosomal and sex-linked components shows that the genetic architecture is highly spatially heterogeneous. For example, orange color in the top of the tail fin and black color ventrally of the dorsal fin were strongly X-linked, while both colors near the pectoral fin showed a large Y-linked component. This overlapping patchwork of genetic architecture generates the complexity of the color pattern.

**Figure 3:**
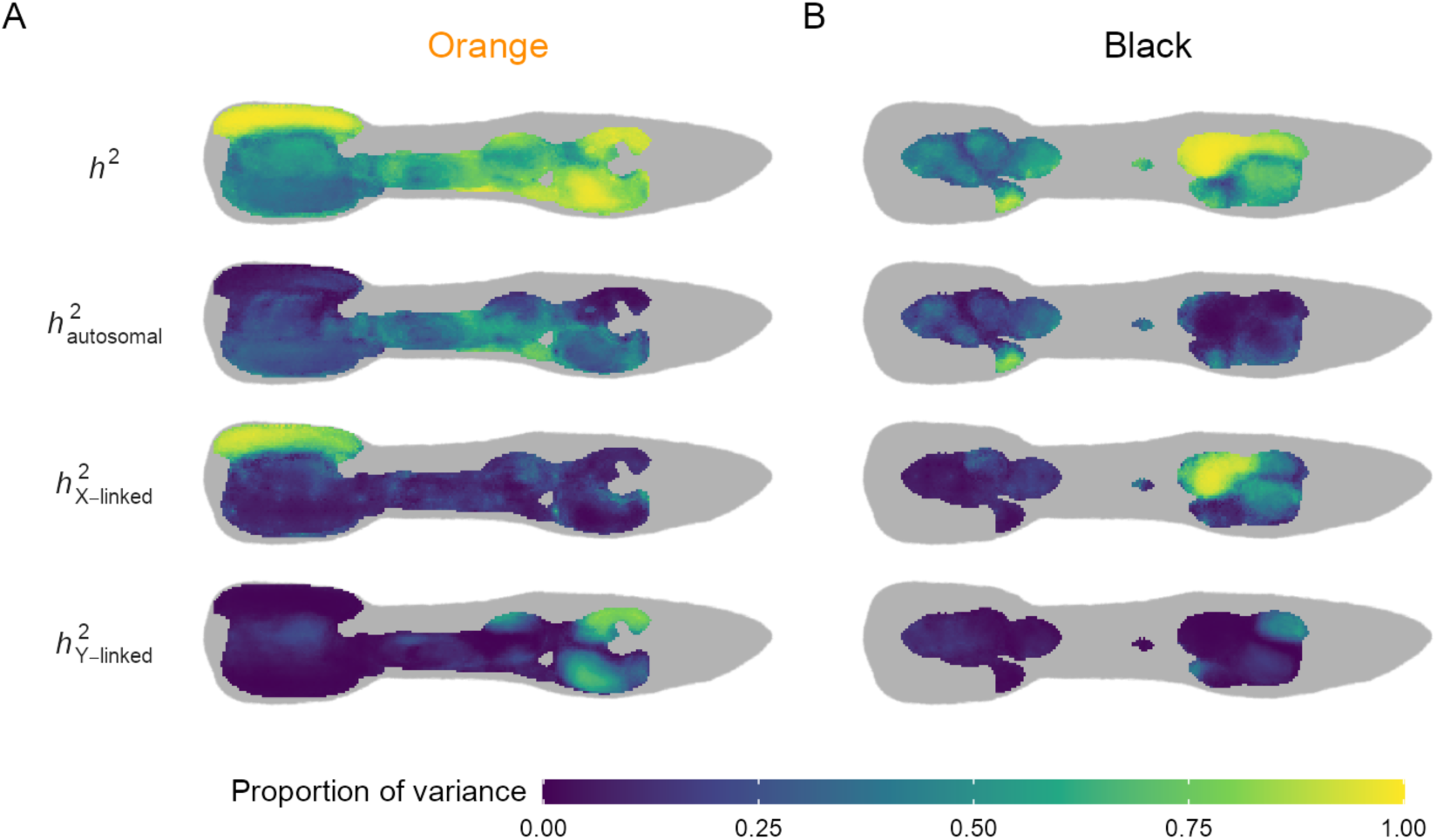
The localized heritability of color is high, but the architecture is heterogeneous. The narrow-sense heritability of the presence of orange (A) and black (B) color on the guppy body. For each heatmap cell, a Bayesian animal model was used to estimate the contribution of additive genetic and environmental factors. Heritabilities are expressed as a proportion of the between-individual phenotypic variance. The autosomal, X-linked, and Y-linked heritability add up to the total heritability. Body positions where the incidence of color is less than 1% are colored grey.

We defined seven orange (Fig 4) and eight black ornaments (Fig S9), based on the incidence of color (Fig 1C) and the localized genetic architecture (Fig 3). As expected, the incidence of all ornaments is heritable. In addition, their size is also heritable for all but one ornament (O3). All orange ornaments responded to selection on the total amount of orange, either in their incidence, their size or both (Fig 4C & 4D). The large effect sizes for ornament incidence are the result of high heritability and strong selection and are consistent with the expectation arising from the breeder’s equation (Fig S10).

**Figure 4:**
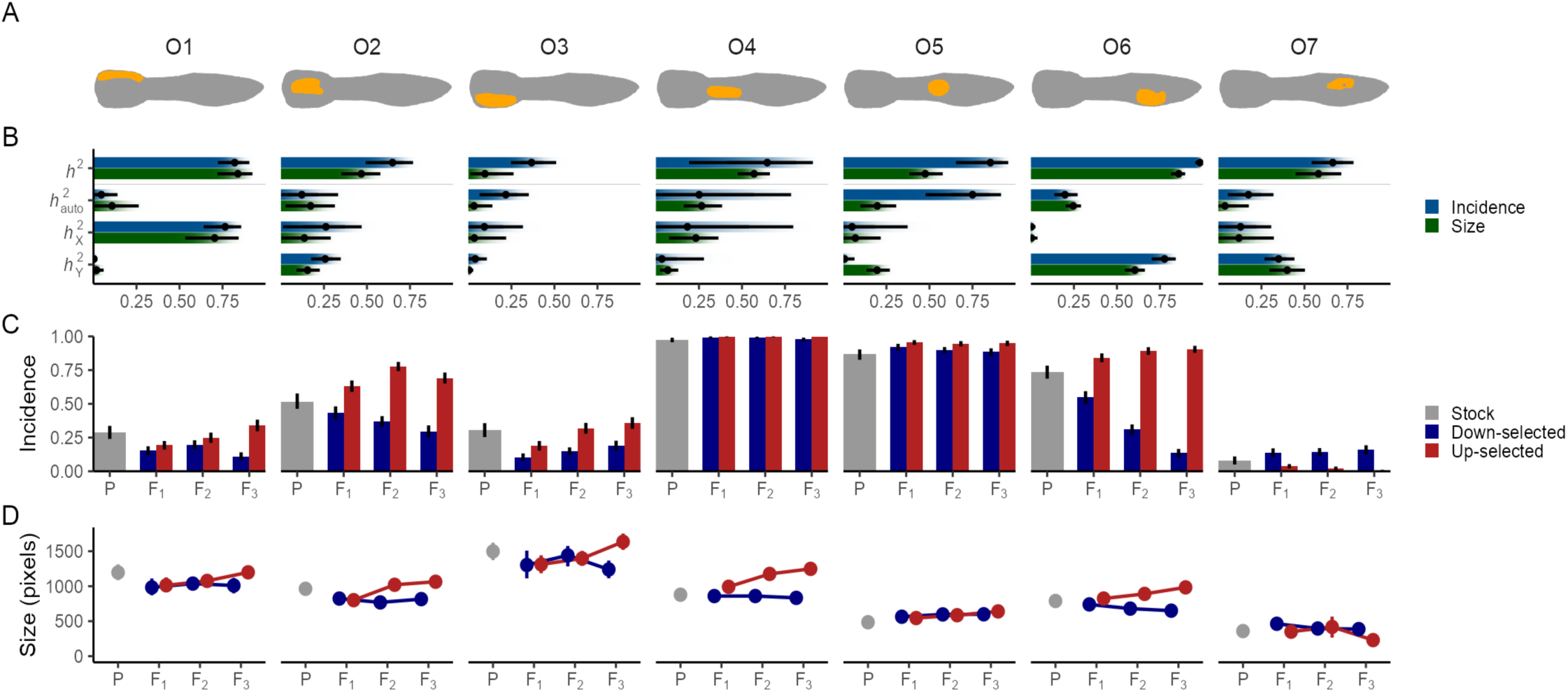
Orange ornaments are strongly heritable and their incidence and size responded to selection. A) Pictograms of seven orange ornaments. B) Heritabilities of the incidence and size (when present) of each ornament. Dots and lines reflect point estimates and 95% credible intervals, and the gradient bars show the cumulative posterior density. C&D) Effect of selection on the incidence and size of orange ornaments. Bars and points represent means, with error bars showing 95% bootstrapped confidence intervals. X-axes show consecutive generations.

Concordant with the localized analysis of inheritance, we find that the ornaments each have their own characteristic combination of autosomal, sex-linked and environmental causes. This can also be seen from the low correlations between many different ornaments with autosomal or X-linked architectures (Figs S11 & S12). In contrast, we observed strong Y-linked genetic correlations, consistent with alleles across multiple loci linked together on a non-recombining Y chromosome. Within each ornament, however, the architecture of incidence and size were typically similar, which may indicate that additive genetic factors for ornament size also control its presence as a threshold trait.

In principle, the seven orange and eight black ornaments can combine into 32,768 (2^15^) unique pattern combinations. We observed 691 of these unique patterns among the 3,229 males in our dataset, with just 130 males (4%) sharing the most common pattern (O2, O4, O5, O6, & B7, Fig S13). Such pattern diversity is consistent with previous qualitative descriptions^8^, although much greater than the only previously reported count of twenty-seven patterns in a different population^5^. We observed fewer patterns than expected from the prevalence of each ornament (1,010), suggesting that linkage disequilibrium between ornaments caused by physical linkage and correlated selection reduced pattern diversity by 32% (confidence interval: [29%, 34%], Fig S14).

These localized approaches will underestimate the true pattern diversity in the population if separate color elements are co-located. For example, some males exhibit large patches of color in the tail that do not neatly fit the three ornaments defined there (Fig S15). Similarly, they do not capture potential heritable differences in spot shape or color characteristics such as hue and saturation. To describe the orange and black patterns free from assumptions about developmental mechanism, we developed a novel high-dimensional phenomics approach using a deep neural network embedder which places the patterns within five-dimensional patternspace. Briefly, using triplet learning^32^, these convolutional neural nets were trained to encode the heritable pattern variation of orange and black patterns (see Supplemental Methods). The networks are only given color patterns but are rewarded when they place patterns from related males closer together than those of unrelated males. As a result, patternspace is a holistic summary of the complex pattern variation among males (Fig S16). Patterns which are close in patternspace share heritable features, as can be observed when we map the presence and size of the ornaments onto this space (Fig S17).

Orange patternspace is divided into two large clusters, differentiated by the presence of the highly heritable and Y-linked O6 ornament (Fig 5 & S17). However, because the Y does not recombine, complete Y-linkage of all color patterns would result in discrete patterns associated with each patriline, which would be expected to form additional clusters, which we did not observe. Instead, the many heritable features combine into a continuous space of pattern variation consistent with high levels of autosomal and X-linked variation across the recombining portion of the genome. Our selection regime gradually separated the populations in patternspace over the generations (Fig 2C). Nonetheless, as there are many routes to increase or reduce the amount of orange, pattern variation is largely maintained within the selection lines among succeeding generations.

**Figure 5:**
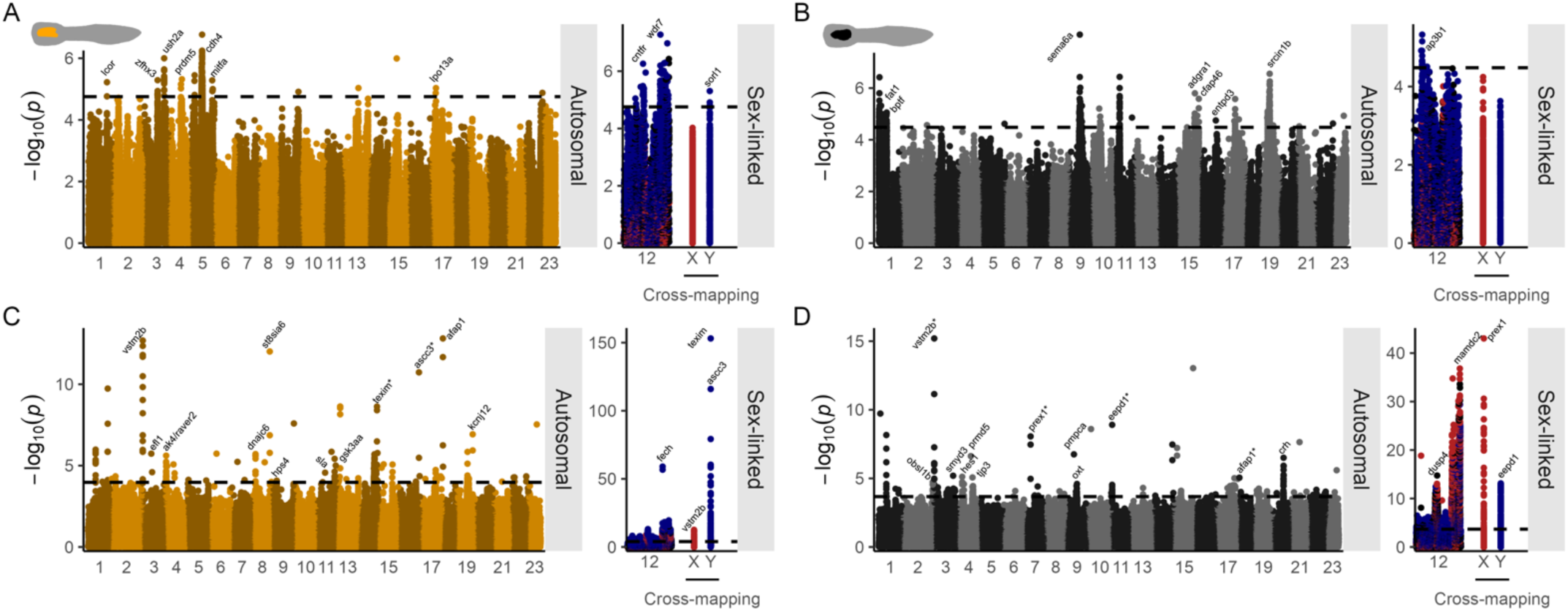
The genetic architecture of color pattern is a mix of autosomal loci and sex-linked loci. A) Manhattan plot showing the association of genomic variants with ornament O2. Points represent genomic variants, with their genomic location on the x-axis, and the p-value of the association on the y-axis. Dotted line denotes 5% FDR. Sex-linked loci are placed to the side, with both the sex-chromosome (LG12) and sex-linked loci cross-mapping to the autosomes shown, colored by their putative origin (red = X, blue = Y). Peaks are labelled with gene names, but unnamed and uncharacterized genes are shown. Names with asterisk (*) indicate that more significant cross-mapping SNPs were present in the same peak. B C, D) As in A, but for ornament B1, orange patternspace, and black patternspace respectively. Results for other ornaments in Figs S19 & S20.

### Mapping color pattern to the genome

We selected 300 males from the F_3_ generation for short-read whole genome resequencing, sampling across all families and across patternspace to maximize the variety of patterns captured. The guppy Y-chromosome is highly diverse, both within^33^ and among populations^34^, and using a male reference genome with a single Y haplotype creates major potential reference bias for all other Y haplotypes (see *Supplemental Methods*). However, the high degree of similarity between the X and Y for homologous regions^33,35,36^ ensures that the vast majority of Y-linked reads will map to the X assembly. We therefore aligned our reads to the guppy female reference genome^37^, which avoids reference bias to any particular Y haplotype, but confirmed our conclusions using a male reference^38^ (Fig S18). After variant calling and filtering, we performed a GWAS for the presence/absence of common orange and black ornaments, and multivariate GWAS for orange and black patternspace. As some of our phenotypes showed sex-linked inheritance, we controlled our GWAS for relatedness across the autosomes as well as the sex-chromosomes (see *Methods*).

The GWAS for ornaments with substantial autosomal inheritance revealed genetic architectures consistent with complex traits (Figs 5, S19 & S20). We recovered multiple peaks on autosomes for each of these ornaments (Fig S21), each explaining a modest proportion of variation in the phenotype (logistic regression, Nagelkerke’s R^2^: <16%), suggesting an oligogenic or polygenic architecture, consistent with threshold trait. To investigate the specificity of these loci, we performed PheWAS and related the genotype at the loci to pigmentation across the body. The loci showed specific local effects for its associated ornament, supporting the conclusion that these ornaments have an independent genetic basis (Fig S21).

Genes with significant variants include those related to cell-to-cell adhesion, which plays a crucial role in the migration of neural crest cells^39^ (NCC), which are the precursors of xanthophores and melanophores^15^. Significant variants include cadherin genes *fat1a, cdh4* and *cdh12* which encode adhesion molecules. Expression of *cdh12* has been implicated in variation in human skin pigmentation^40^, and other cadherins have been associated with population-level differences in guppy coloration^41^. In addition, a variant close to *sema6a* explained 16% of the variation in the tail ornament B1. This gene is a target of the melanocyte master regulator *mitf* in humans^42^, is upregulated during the growth of pigmented tail fins in betta fish^43^, and semaphorins have been broadly implicated in guiding NCC migration in zebrafish^44^. Similarly, *sema4f* is associated with ornament B6. Chromatin remodeler *bptf* is necessary for *mitf* function^45^, and is also implicated in ornament B1. We also find signal upstream of *mitfa* itself for ornament O2, supporting a recent study showing *mitf* also affecting the fate of xanthophores in medaka and zebrafish^46^. A variant associated with B6 falls in *lin28a*, a key regulator of position-dependent NCC differentiation^47^. *Cygb1* affects NCC development and melanophore patterning in zebrafish^48^, and *fscn1a* is involved in directional NCC migration^49^. Together, these results suggest that many of the ornaments are ultimately under the control of loci modifying the trajectory of NCCs, as they migrate towards the skin and develop into chromatophores.

Next, we investigated the genetic basis of color traits with X- and Y-linked patterns of inheritance. For orange and black patternspace, highly significantly-associated variants span the entirety of linkage group (LG) 12, the sex chromosomes, largely due to strong linkage across this region. Given the low differentiation between the X and Y for homologous regions^33^ (male:female coverage at LG12: 21-26 Mb = 0.986), most reads from the Y chromosome map to the X chromosome in the female reference genome, and variants on LG12 represent a combination of pseudo-autosomal, X-linked and Y-linked variants^35^. Ornament level GWAS (Figs S19 & S20) confirmed that ornaments with strongly sex-linked inheritance are associated most with LG12.

Surprisingly, we observed additional peaks of associated variants on nearly all autosomes for ornaments that our heritability analysis showed to be sex-linked. Strikingly, many of these regions showed excess heterozygosity, elevated read depth, and an increased read depth from heterozygotes compared to homozygotes, suggesting reads from copy number variants (CNVs, Fig S22). Given recent work that identified CNVs spanning autosomes and the Y chromosome in natural guppy populations^50^ and the role of sex-linked variation in this system, we hypothesized that reads from the Y chromosome regions that are not homologous to the X could map to these autosomal regions, representing CNVs spanning multiple genomic locations. We tested this conjecture by comparing the genotypes at each variant with expectations from autosomal, X- or Y-linked segregation. Briefly, we modelled the dose of the variant as following the pedigree-based covariance between individuals in each of the three scenarios. We then called the putative source of these variants as X- or Y-linked if their relative likelihood (AIC weight) exceeded 80% (Fig6). Using this pedigree-informed approach, we identified that 1% of called variants were potentially the result of cross-mapping reads between the autosome and sex-chromosome. Variants implicated in the patternspace GWAS were highly enriched for these cross-mapping variants (orange pattern: 19 X-linked & 154 Y-linked out of 510 called variants; black pattern: 46 & 252 out of 563; Figs S23 & S24), also suggesting the involvement of CNVs with sex-linked copies.

One possible explanation for the abundance of Y-linked variants in our GWAS is that they are all strongly linked to one or a few causal loci of large effect which control the Y-linked components of the color traits. To test this, we performed principal component analysis on the variants associated with orange pattern and a Y-linked inheritance pattern. If these loci have significant associations with orange pattern due to strong linkage to the same few alleles, we expect these variants to cluster into clear haplotypes that are strongly associated to color. Instead, we identify four putative Y-haplogroups with substantial variation within each haplogroup (Fig S27), and which only explain a small amount of color variation compared to individual variants (Figs 6C & S28). The fully non-recombining region of the Y chromosome is relatively small^36^, and the majority of the chromosome recombines with the X exceedingly rarely^51^. Although these rare X-Y recombination events might explain the variation without our four Y-haplogroups, this would also lead to expected X and Y segregation of SNPs. We therefore suggest that our data are more consistent with a substantial amount of Y diversity in our population.

**Figure 6:**
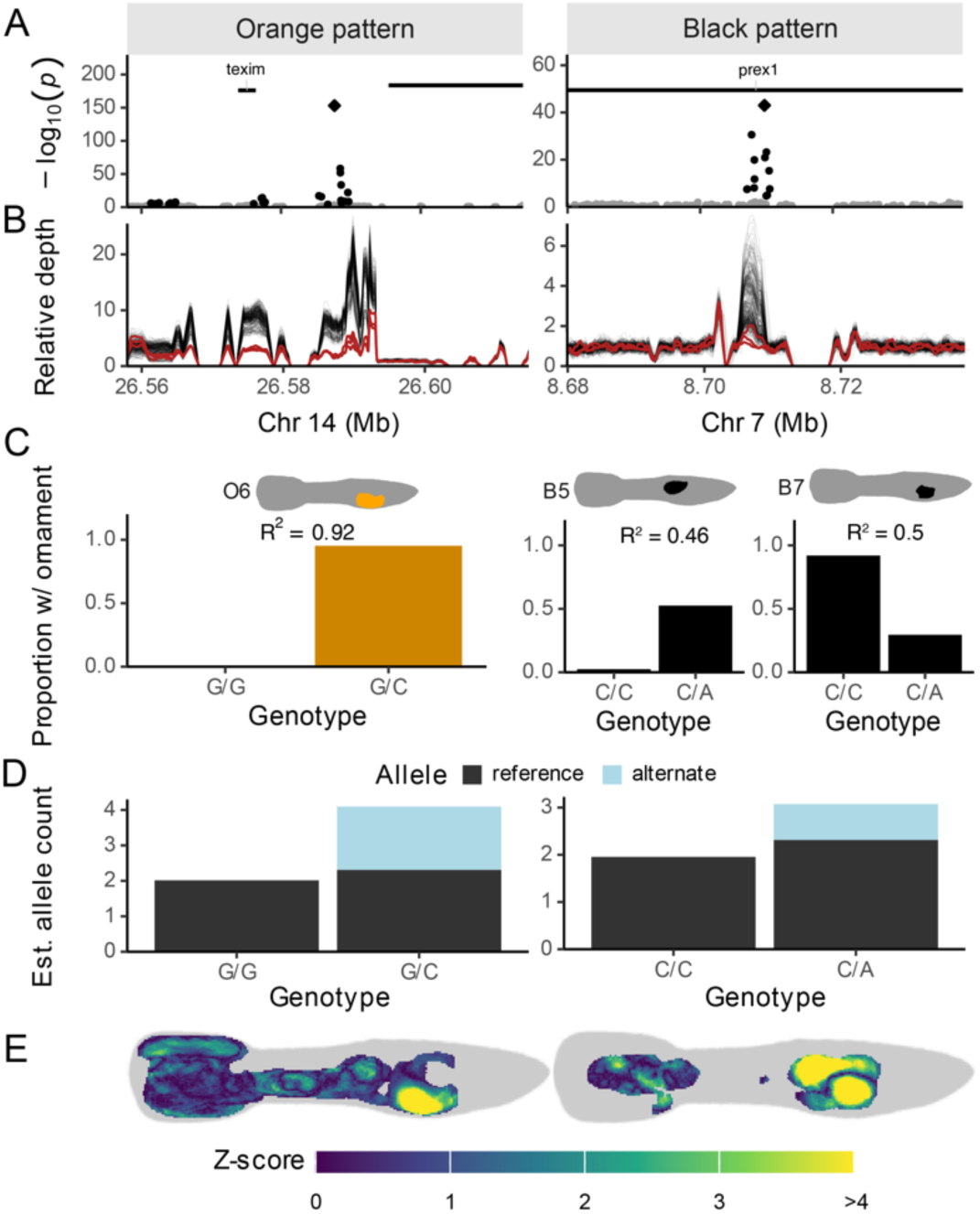
Large effect sex-linked CNVs underlie sex-linked variation in color pattern. A) GWAS peaks on autosomes with significant loci with sex-linked inheritance, near *texim* and in *prex1*. B) Relative coverage (1x = genome wide sample average) across the region for 297 males (this study) and 3 females, calculated in moving windows of 1kb, 100bp apart. C) Phenotypic effects of the top variant (diamond in A) on ornaments O6 (orange pattern) and B5 & B7 (black pattern). The proportion of variation in the phenotype that is explained by the top SNP is displayed as Nagelkerke’s R^2^ from a logistic regression. D) Median estimated allele counts for the top SNP, based on read depth, for homozygous reference and heterozygous individuals. No heterozygous alternate individuals were found. E) PheWAS heatmaps displaying Z-scores for the effect of the top variant on orange and black color across the body, controlling for autosomal and sex-linked relatedness.

The locus most strongly associated with orange pattern mapped to LG14 (an autosome), with many significant variants in a 35kb region that are inherited in a Y-linked manner (Fig 6A, left column), suggesting these variants represent fixed differences between the autosomal region and its copies on the Y-chromosome. Consistent with this scenario, we observed highly elevated coverage consistent with multiple Y-linked copies in males, and reduced coverage in females (Fig 6B).

The gene located in this region (LOC103476393) is uncharacterized in the annotation but has 72% sequence identity with *teximYa* from the related Southern platyfish (*Xiphophorus maculatus*)^52^. Y-specific duplication of *texim* has been documented in multiple guppy populations^38^, and copies of *texim* have also been found on the Y-chromosome of the closely related *Poecilia parae*^53^, suggesting duplication of *texim* to the Y-chromosome is common in this clade, and most likely mediated by helitrons^38,52^. In fact, the most strongly associated variant in this region is located downstream of this *texim* copy, and falls in the *texim*-associated helitron. This top SNP predicts 92% of the phenotypic variation of the Y-linked ornament O6 (Fig 6C & 6E). Coverage for most of this region is elevated five to ten-fold and heterozygous individuals have increased coverage compared to individuals that are homozygous reference (Fig 6D), so that coverage in this region is strongly predictive of the presence of ornament O6 (Fig S29). Using long-read RNA sequencing, we confirmed the existence of at least five *texim* copies whose presence is polymorphic on the Y-chromosome. We also confirmed that the helitron is transcribed together with the *texim* gene. One such copy is expressed in adult tail tissue, but only in males, again consistent with Y-linkage (male vs. female: 2.14 vs 0.00 CPM; t_4_ = 3.8, *p* = 0.019; Fig S30). Genotyping the males in our GWAS for the presence of this copy revealed that it is predictive of the presence O6 (R^2^ = 0.30, Fig S29), but can only partially explain the observed association of the top GWAS SNP (R^2^ = 0.92).

Similarly, the locus most strongly associated with black pattern mapped to LG7, also an autosome (Fig 6, right column). We observed many highly associated sex-linked variants in a 3.5kb region, again paired with an increased read-depth for males, but not females. This locus all shows non-Mendelian inheritance with increased read depth for heterozygotes and a lack of alternate homozygous genotypes, all supporting the notion that these variants are the result of cross-mapping from the sex-chromosomes. The inheritance pattern of the significant SNPs indicate that the presence or absence of an X-linked copy is most strongly associated with the phenotype, although increased coverage in males compared to females suggests the presence of additional Y-linked copies. All significant variants fall into the gene PIP_3_-dependent Rac-exchanger 1 (*prex1*). This gene has previously been found to be important to melanoblast migration in mice, where disruption of this gene leads to white paws and belly, the most distal endpoints of NCC migration^54^. This locus alone explains 46% and 50% of the phenotypic variation in ornaments B5 and B7, but in opposite directions. Since these ornaments are in close vicinity, this may suggest that genetic variation in *prex1* modifies NCC trajectories to terminate in one or the other location. Overall, the sex-linked ornaments appear to have a monogenic or possibly oligogenic architecture, with evidence suggesting a large role of copy number variants that span the autosomes and sex chromosomes.

## Discussion

The remarkable variation in male guppy coloration has made them a model for studies of female preference^5,6^ and the interaction of sexual and natural selection^4,55^. This variation has long been suggested to have a simple genetic basis, but the large variation in phenotypes has precluded analysis of the full complexity of the pattern and identification of responsible genomic loci. Here we used a combination of high-resolution phenotyping via neural nets, controlled pedigrees, and GWAS to dissect the detailed genetic architecture for male color variation.

Our population has shown a rapid response to artificial selection for other complex traits^56–58^, suggesting significant underlying genetic variation. Consistent with this, we observe substantial variation in male color (Fig 1) and a rapid response to artificial selection for increased and decreased orange coloration in all our replicate selection lines (Figs 2, 4). Male color has previously been shown to respond to artificial selection in other populations^10,59,60^, and consistent with this, we observe high heritability (Figs 3 and 4) for both total color area and specific color ornaments. Nonetheless, we see that color is not entirely genetically determined, and there is a significant role of the environment for the size of ornaments and locations in the tail fin. The role of color as an honest signal of a male’s ability to e.g. forage and fend off parasites^1,2^ may either be mediated through these less-heritable components of the color phenotype, or color loci may be genetically linked to loci conferring such fitness benefits.

We confirmed that sex-linked variation plays a substantial role in the genetic architecture of color in this population, although for only a subset of ornaments (Fig 3). Previous reports on other populations based on the visual assessment of select co-inherited ornaments have often emphasized their Y-linkage, although there are also many which have been observed segregating on the X or autosomes^5,21,24–26,61^. It is therefore unclear how much of the total color variation these classic patterns capture. Quantitative analyses based on color area and spot number have also shown a larger contribution of the sire genotype, but dam heritability nonetheless plays a significant role^11,62^. Finally, testosterone-treated females will generally express some ornaments, demonstrating that complete Y-linkage is the exception^16,26^. As both the observable ornaments and their sex-linkage varies considerably among populations, we look forward to population comparisons using the comprehensive analyses now available.

One might expect that if black is epistatic to orange, selection to increase the area of the fish with orange coloration would reduce the total black area. That is not the case, as we observe little change in black area throughout the selection experiment but do see modest localized shifts in black color (Fig S3), more consistent with the repulsion of developing xanthophores and melanophores during color development^63^.

We then used whole genome sequencing for 300 males from the F_3_ generation, sampled to maximize their distribution within color-space to perform GWAS for multiple color traits (Figs 5 and 6). Interestingly, genes previously identified as key to melanophore formation in guppies, *colony-stimulating factor 1 receptor a* (also called *fms*), *kit ligand a* (*kita*)^27,64^ and *adenylate cyclase 5* (*adcy5*)^65^, were not implicated in our GWAS (Fig S31). This suggests that although these loci are fundamental and necessary in the development of color cells, they do not contribute substantially to the variation in color pattern we observe in our population. Instead, we find significant associations with genes involved in cell-cell adhesion and neural crest cell migration, suggesting that the migration routes of chromatophores and their pre-cursors during embryonic development, rather than the production of pigments, determines the black pattern. Genetic variation in one such gene, *prex1*, explains half the phenotypic variation in two black ornaments.

In contrast to melanin, carotenoids are not synthesized in vertebrates *de novo*, but rather collected from dietary intake and modified prior to deposition^66^. This mode of color development is hypothesized to make carotenoid-based coloration a key mediator of honest sexual signaling of male quality^1,67,68^. Consistent with this, males from selection lines with greater orange area exhibited greater sexual vigor (Fig S4 & S5), though we did not observe any correlated changes in female life history traits. The behavioral response to selection on color may be the result of pleiotropic changes to neural crest cell development, which plays a central role in both pigmentation and neurological development^27,28^.

We might expect genes associated with carotenoid metabolism in other vertebrates to play a role in orange coloration in guppies. A key gene identified in red coloration of birds, cytochrome P450 locus CYP2J19^69,70^, lacks a reciprocal ortholog in teleost fish. Aldehyde dehydrogenase family 1 member A1, associated with red carotenoid coloration in lizards^71^, is also not implicated in our GWAS. Instead, the region most strongly associated with orange color is a helitron that is closely linked to and co-expressed with *texim*. Although the functional role of *texim* remains unclear, it is thought to encode an esterase/lipase^52^, important in carotenoid modification^72,73^ and therefore may play a role in orange coloration.

Similarly, we do not find support for a large haplotype on LG1, which has been previously been indirectly implicated in guppy color variation in another population^74^. Instead, our results suggest a far more complicated genetic architecture for color pattern. For both black and orange color, we observe specific ornaments encoded through largely independent suites of loci (Figs S23 & S24). Most ornaments have a polygenic and largely autosomal architecture. Moreover, the sex-linked architecture is heavily influenced by CNVs, although it is challenging to differentiate between gene dose effects, with more copies of a gene associated with greater color, and allelic variation, with some copies associated with specific patterns. CNV dose has also been implicated in rapidly evolving color phenotypes during domestication. In rock pigeons, at least seven alleles that vary in the copy number of the Z-linked *st* locus produce different depigmented phenotypes^75^, while additional copies of a CNV near the melanin-associated *kitlg* gene causes darker coats both within and across dog breeds^76^. This aspect of the genetic architecture of guppy color is particularly interesting in that it might be expected to help maintain male phenotypic color variation more effectively than allelic variation in single-copy genes. It has been a long-standing mystery how the remarkable variation in male color is maintained against the eroding forces of genetic drift, selection and sweeps. Although female mate preferences for rare male patterns can be expected to preserve some male patterning variation^5,13,14^, it is difficult to envisage how this would be sufficient to maintain the extensive variation observed in guppy populations, many of which formed with significant bottlenecks and founder effects. Moreover, the evolution of female preference for rare patterns has been proposed to be beneficial to females through the indirect benefit of sexy sons^5,77^. However, such benefits rely on a strong genetic covariance between the pattern of fathers and their sons. It is unlikely that populations like the one studied here, with its many autosomal and X-linked inherited pattern elements, would confer such benefits to females.

The fact that we observe CNVs associated with color spanning autosomes and the sex chromosome may help to explain recent observations of variation in the extent of Y-linkage for color in wild populations^16,22^. Although the size of the fully non-recombining region of the Y chromosome is relatively small^34–36^, there is evidence that it can accumulate substantial recent duplications of genes from the remainder of the genome^50^. If this process involves coloration loci and varies across populations, it is easy to envisage how the association between the Y chromosome and color variation could differ, even among closely related populations. It may also suggest that the comparatively low-levels of Y-linkage we observe in our population, originally collected from the high predation region of the Quare River, Trinidad, may be observed in other natural populations that either have lower rates of duplication to the Y, or lower levels of Y-duplicate retention. Finally, our data suggests that the Y-linked regions that do contribute to color in our population are surprisingly diverse.

Taken together, our results present a critical link between the extraordinary variation in male guppy coloration and the genome that encodes it. The fine-scale genetic architecture of color that we reveal presents an exciting opportunity for future molecular analyses of sexual selection dynamics and a critical point of comparison for future studies of the genetic architecture of highly polymorphic traits.

## Acknowledgements

We thank Lengxob Yong for sharing insights on guppy photography. We thank Ben Sandkam, Ben Furman, David Metzger, Lydia Fong, Yuying Lin, Christina Hodson, and James Lewis for help with animal husbandry and valuable discussions, and Tom Booker for advice and comments on an earlier draft. Yuying Lin contributed the female coverage data. Funding was provided by a Canada 150 Research Chair, NSERC and an ERC grant (680951) to JEM.

## Author contributions

Conceptualization: WvdB, AC-L & JEM, Methodology: WvdB, Formal analysis: WvdB & AC-L, Investigation: WvdB, JJS, VSG, LMS, CJ, MC-C, AC-L & JEM, Writing – Original Draft: WvdB & JEM, Writing – Review & Editing: WvdB, JJS, VSG, LMS, CJ, MC-C, AC-L & JEM, Visualization: WvdB, Supervision: JEM, Project administration: JJS, Funding acquisition: JEM.

## Data and code availability

Code is available at https://github.com/Ax3man/vdBijl_etal_2024_GuppyColorPatterns. Sequence data will be deposited upon publication.

## Materials and methods

### Study population and animal husbandry

Our study population are descendants of guppies that were captured in Trinidad in 1998 and subsequently maintained in large, outbreeding stock populations to maintain genetic diversity, and males show large diversity in their ornamentation. 240 females for the parental generation (P) were randomly sampled as juveniles from our three stock populations and maintained as virgins until sexual maturity. 300 males for the parental generation were sampled as adults from the stock populations.

Fish were kept in 1.4 l tanks in three flow-through systems, with each experimental replicate in a separate system. Water was maintained at 25 °C, and lights were on a 12:12h cycle. Fish were fed daily with a mix of live brine shrimp and tropical fish micro pellets (Hikari, Japan), with immature fish fed an additional meal on weekdays of frozen daphnia and bloodworms.

### Experimental procedure and selection

Each replicate was started using 100 males, evenly sampled across the three stock populations. We quantified the amount of orange coloration for each male (see *Color pattern extraction and quantification*, below), and selected the 30 males with the largest and 30 males with the smallest total orange area to form paired up- and down-selected lines. For the P generation, these males were then randomly paired with females. To increase our statistical power to detect sex-linked variation, we paired 10 males of each line with two females each to generate half siblings, and 20 males with a single female each, creating a partial half-sib breeding design. For subsequent generations, we selected the 30 males with greatest orange area in up-selected lines, and the 30 males with least orange for down-selected lines. Additionally, we increased the strength of selection by selecting on females. Since females do not express orange coloration, we used an animal model to estimate breeding values (see *Quantitative genetics,* below). We then selected the females with greatest and least breeding value (40 for each category), incorporating uncertainty by ranking across draws from the posterior distribution. To avoid a rapid loss of genetic variation, and to limit the selective advantage of males paired with two females, we further limited our selection to include a maximum of three sons from each sire, and three daughters from each dam.

For the males matched with two females, females were kept in separate tanks, and the male was moved between females each week. We collected broods from the up- and down-selected lines until we had 90 broods per generation per replicate. 93.3% of paired males and 89.3% of paired females successfully sired offspring, resulting in an average of 1.46 broods per sire, and 1.10 broods per dam. Having >1 brood per pair allowed us to estimate the magnitude of brood effects.

Broods were collected within a few days of birth to avoid infanticide and moved to their own tank. To maintain similar developmental rates, broods larger than ten individuals were randomly reduced to ten. Broods were checked three times per week for signs of male maturation (first signs of gonopodium growth), and males were moved to a separate tank to avoid sibling mating. The tanks with male brood members were then queued for phenotyping after visual confirmation of sexual maturity (fully developed gonopodium and color pattern), and a minimum age of 60 days. After phenotyping (see *Color pattern extraction and quantification*, below), males were housed in individual tanks to maintain their identities.

The three replicates were set up staggered in time, so that one replicate could be phenotyped while the others were breeding and maturing. Fish were only marked with their individual IDs, blinding the experimenters as to which line individuals belonged to throughout experimental procedures and phenotyping (except when setting up the next generation of breeding pairs).

### Animal photography and calibration

We (WvdB & JJS) captured photos of male color patterns using a Canon EOS Rebel T7i (Canon, Japan) with Canon EF 100mm f/2.8 Macro USM lens (Canon, Japan). All photos were captured using identical camera settings, at an aperture of f/11, an exposure time of 1/320s, an ISO of 800 and an approximate distance of 35-40 cm, and we captured a color standard *in situ* before each photography session (ColorChecker Passport Photo 2, X-rite, USA). Photos were captured in RAW format, then color corrected in Adobe Lightroom (Adobe, USA), see *Supplemental methods*.

The photarium (photo tank) was placed on a pedestal covered with matte black foam, inside a 30 x 60 x 30 cm photo box. The walls and top of the box were constructed of opaque white plastic, to minimize the influence of external lights. In the top, directly above the pedestal, we mounted 2 Neewer NL660 LED panels with white diffusers, with a color rendering index of >= 96 (Neewer, China).

We photographed both sides of every fish, and for 99.9% of fish we had at least three photos, with the replicate photos allowing us to parse measurement error from asymmetry. We analyzed a total of 14,100 photos, for an average of 4.37 photos per fish.

### Color pattern extraction and quantification

After color correction, we used a custom pipeline to analyze the images. The pipeline consists of deep neural networks for image segmentation and landmark placement, and geometric morphometrics for image alignment. To be able to phenotype fish during the selection procedure, when we still had little data available to train the models, we adopted a semi-autonomous strategy. We used an interactive script to phenotype fish, which prompted the experimenter (WvdB) to visually check the output of each analysis step, and correct manually when the model prediction was incorrect. We then intermittently updated model weights with the growing number of manually annotated examples to improve model performance, gradually reducing the need for manual intervention throughout the experiment. The pipeline consisted of five steps (SI). By comparing the number of pixels in the extracted fish mask from Step 1, and the number of pixels in the extracted carotenoid and melanin masks from Step 2 (i.e. before warping), we calculated the amount of coloration as a percentage of the body area. For use in our selection paradigm, we calculated the mean across multiple photos of each side of each fish, and then the mean across the sides.

### Quantifying effects of selection

We quantified the effect of selection on the amount of orange and black coloration using Bayesian linear mixed models (LMMs) using *brms*^78,79^. We modeled the amount of color as a percentage of the body area as the dependent variable, with fixed effects of generation, selection direction and their interaction, and a random intercept per generation nested within experimental replicate. We then calculated the contrast in marginal means between up- and down-selected fish of the F_3_ generation using *emmeans*^80^. We used the same modelling approach for the effect of selection on life history parameters, except that we modelled fecundity using the negative binomial distribution. In addition, we estimated the effect of selection on orange and black color per body location (Fig 2B & S3) using independent binomial generalized linear mixed models with *lme4*^81^, using the same model structure, and the presence/absence of orange as the dependent variable.

### Male sexual behavior

We assessed the sexual behavior of 126 male offspring from the F3 generation (63 from each selection direction, evenly split across the replicate lines). We isolated males upon early indications of sexual maturity (first signs of gonopodial development), then at around three months of age tested their sexual behaviors using a no choice test with nonreceptive females, a standard for studying male sexual behavior in poecilids^30^. Behavioral trials took place in a 47 cm diameter arena, with fish transfer followed by a 5-minute acclimation to minimize stress and a 15-minute observation using a webcam at 1080p and 30 fps. We analyzed the recorded videos to quantify male sexual behaviors. We utilized idTracker^82^ for positional data during each trial. Trials where females exhibited receptivity or where fish showed signs of stress were excluded, leading to a final dataset of 95 individuals. Statistical analysis on behavioral differences between up- and down-selected males used generalized linear mixed models implemented in *glmmTMB*^83^, with behavioral metrics as dependent variables, selection direction, male body size, and female body size as fixed effects, and replicate line as random effect. Behavioral counts were modelled using the negative binomial distribution, while for time spent following we used the Gaussian distribution with log link function.

### Ornaments

We defined areas of the fish as ornaments manually, based on the visible spatial structure in the incidence of color and the localized genetic architecture (Figs 1C & 3). We then overlayed these areas on the extracted color patterns from each photo and counted the number of pigmented pixels within each area. We scored an ornament as “absent” (ornament size = 0) if < 10% of the ornament area was pigmented. This limit was chosen based on the bimodal distribution of the number of pixels, as in some cases a few pixels from a neighboring ornament spilled over into other areas.

### Quantitative genetics

Estimating of quantitative genetic (QG) parameters for the amount of orange and black color was performed by MCMC sampling of Gaussian animal models implemented in *Stan*^84^ with *brms*^78,79^. Additive genetic variance-covariance matrices were calculated from the pedigree using *nadiv*^85^, with the *S* matrix for X-linked effects assuming no global dosage compensation^36^. Y-linked genetic variances were estimated by including patriline as a group level effect. Similarly, we included dam identity to estimate maternal effects, included brood identity to capture shared environmental effects between siblings, and individual identity to estimate the remaining environmental influence, all as group level effects. We included the side of the fish, nested in individual identity, as a group level effect to estimate the variance due to asymmetry, and included the residual variance to estimate measurement error. For all variances, including the residual variances, we estimated their correlation among the two colors. We used default non-informative priors for all parameters.

For the QG heatmaps (Fig 3), we estimated separate animal models for each cell, estimating orange and black colors separately. We used the same model structure as the model above, now with a Bernoulli error structure. We also removed the maternal effect, as the clutch and maternal effects are correlated and their magnitudes are small. Since estimating these parameters independently thousands of times using MCMC is prohibitively computationally expensive, we used an approach that makes use of the fact that the QG estimations will be similar between neighboring cells. To limit the number of models, we reduced the dimensions of the color pattern images by a factor of two (250 x 70 cells or pixels), and only ran models for cells with an incidence of at least 1%. We then performed model fitting in two rounds.

First, we fit animal models for the cells with x- and y-coordinates divisible by five. For these models, we used four chains, with 500 warmup iterations and 1500 sampling iterations per chain, and uninformative StudentT(3, 0, 5) priors on the intercept and variance components. In a second step, we then used the output of these models to fill in the cells between this five-by-five grid. These models only used two chains of 500 warmup iterations and 500 sampling iterations, but were seeded with initial parameter values and step sizes that were the average of the closest nine models from Step 1, weighted by their inverse distance. Similarly, we set normal priors for the intercept and variance components with the mean (also weighted by inverse distance) and standard deviation of those parameters of the closest nine models.

For ornament heritability, we used similar models to the amount of orange and black color, but used hurdle-lognormal models instead to jointly estimate the variance components for the incidence and size (when present) of the ornaments. Given the very small maternal effects on total color, we did not include that parameter here. We specified an uninformative logistic(0, 5) prior on the hurdle parameter, and a StudentT(3, 0, 5) parameter on the standard deviations of the group level effects, with otherwise default priors.

For the pairwise correlations across ornaments, we used the Gaussian approximation and fitted the multivariate animal model on individual-level means using the restricted maximum likelihood method from *sommer*^86^. Given the small magnitude of brood effects from the full Bayesian animal models for each ornament, we did not include that effect here.

### Learning of patternspace

We devised a deep learning approach to map images of guppies onto a “patternspace”. Our objective was to capture the heritable components of color patterns by learning a space where images of related guppies are placed closer in this patternspace compared to those of unrelated males. Our method is predicated on the “triplet loss learning” paradigm^32^. In this paradigm, a model maps images into a space, one at a time, using a learned metric. Because the model uses the same metric for each image, images that are placed close together are similar according to that metric. As our goal was to obtain a space describing the heritable variation in color patterns, we trained the model to use a metric that captured this heritable variation. This training used triplets of images, which comprise an anchor image (a random male), a positive image (of a related male), and a negative image (of an unrelated male). By comparing the distances between these three images, we can evaluate whether the metric is capturing heritable variation. During training, each image is passed through the model and the triplet loss function is applied, optimizing the metric to reduce the distances between anchors and positives, while increasing the distances between anchors and negatives. Unrelated males had a coefficient of relatedness *r* of 0, while for related males *r* was at least 1/8.

The deep learning architecture was designed to process images of dimensions 250x70x3, with the last dimension being color channels. Our model, a Convolutional Neural Network (CNN), had a series of layers that successively compress the images to lower dimensions. Specifically, we used separable convolutional layers with a kernel size of 3x3 and *relu* activation, followed by a max pooling layer, repeating this four times. Each convolutional layer had double the filters of the previous, starting at 8 filters. After the fourth pooling layer, we flattened the output to a dense layer of size 8, and then a second dense layer of the embedding dimension. Lastly, a batch normalization layer normalized activations to unit variance. Embeddings were obtained at this normalization layer.

We trained for 50 epochs using a batch size of 128 images. At each epoch, we randomly generated 5,000 triplets. We randomly allocated 20% of individuals to the validation set, from which we generated 500 triplets per epoch to track model performance. Moreover, learning rate schedules were implemented to adjust the learning rate across epochs, with specified rates of 10^-4^, 10^-5^, and 10^-6^ at epochs 0, 30, and 40 respectively. This scheduling ensured a faster convergence initially, with fine-tuning during the later stages of training.

We evaluated the number of embedding dimensions by training the model at one to eight dimensions, then chose five dimensions, since the model accuracy for the validation sample stopped increasing above five. After training, we used the CNN to embed all 14,100 images. All analyses use the five embedding dimensions while for visualization we performed dimension reduction using UMAP^87^, which aims to preserve the local neighborhood structure of the data, as implemented in R package *uwot*^88^.

### Genetic sampling and whole genome sequencing

To maximize power for the genetic mapping of color traits, we sampled 50 fish from each selection line (N = 300) for whole genome sequencing (WGS). To maximize genetic and phenotypic variation in our sample, and therefore power, we sampled males from each family with offspring in generation F_3_, and sampled males with the most/least carotenoid coloration in up/down-selected lines respectively. This meant we sampled one son from each family, with some families contributing a second son until we reached the desired 50 males. Additional methods are in the SI.

DNA was extracted from sampled males using Qiagen Dneasy blood & tissue kits (Qiagen, USA). Samples were then sequenced on four lanes of Illumina NovaSeq 6000 S4 PE150 to an average depth of 10x per individual. One individual was not included because the library failed. We checked the identity of samples by visually comparing the genome-wide relatedness (GRM) with the pedigree-based relatedness (A) and excluded two samples which did not match, leaving 297 samples for analysis.

### Genome-wide association analysis

We first trimmed the adapters using *Trimmomatic* and then mapped the reads to the reference using *bwa mem* using default mapping parameters. We marked PCR duplicates using GATK’s *MarkDuplicates*. We then called variants using *bcftools mpileup* where we downgraded the mapping quality of reads containing mismatches with *–adjust-MQ 60*, and set a minimal mapping quality using *–min-MQ 30*, followed by variant calling using the consensus caller *bcftools call*. We then performed variant filtering, and dropped sites with a QUAL score < 30, MQ < 40, FS > 60, |BQBZ| > 10, |MQSBZ| > 10 or |RPBZ| > 10, and sites where more than 20% of individuals had a depth < 5, or > 20% of individuals had a GQ score < 10. Additionally, we filtered out sites with a major allele frequency < 10% or a missingness > 5%, as well as indels.

We performed GWAS using the two-step “population parameters previously determined” (P3D) approach^89^ implemented in *sommer*^86^. We controlled for relatedness and population structure by including random effect terms for the autosomal, X-linked and Y-linked portions of the genome, as well as a fixed effect for selection direction. We determined the population level parameters using the full set of phenotyped individuals. For the autosomal relatedness, we combined the pedigree-based matrix *A* with the genomic relatedness matrix (GRM) estimated using GEMMA^90^ for the sequenced samples, to obtain the single-step matrix *H*^91^. We used the *S* matrix for X-linked relatedness, and patriline identity for Y-linked relatedness. We then performed marker-wise effect size estimation for the subset of sequenced individuals. We performed multivariate GWAS by performing this procedure on each of the 5 axes of patternspace. We then combined the Z-scores to obtain multivariate p-values using CPASSOC^92^. For univariate ornament GWAS, we used the same approach, using the normal approximation, and only including ornaments with an incidence between 5% and 95%.

The phenotypic effects of individual variants were evaluated with PheWAS using *sommer*, where we controlled for autosomal, X- and Y-linked inheritance, fixing those variance parameters at the posterior medians of the per-pixel quantitative genetic analyses described above.

Coverage statistics were calculated using *samtools depth*, with a window size of 1000bp and a window distance of 100bp. We additionally calculated coverage for sequencing data from 3 females from the same population taken from^93^ (NCBI BioProject accession PRJNA858015, sequenced at ∼30x coverage), which were re-analyzed using the identical pipeline used for the males from the current study. We controlled the false discovery rate (FDR) using the Benjamini & Hochberg (BH) procedure^94^ separately for the orange and black patternspace GWAS. For ornament GWAS, we performed the BH procedure jointly across the ornaments, but separately per color. FDR is controlled to 5% within each of these four sets of tests. We compared the overlap in associated variants between traits using the *ComplexUpset* package^95^.

We performed principal component analysis to detect Y-haplogroups using *irlba*^96^. We assigned the haplogroups using hierarchical clustering on the Euclidian distance matrix of the first 20 principal components using *hclust*.

### Inference of variant inheritance pattern

The inheritance pattern of all called variants was estimated using animal models. We encoded the genotypes as doses of the alternative allele, and then fitted three models using *sommer*^86^ with dose as the dependent variable. The three models included a random effect for the autosomal, X-linked or Y-linked covariance, respectively, as described in *Quantitative genetics*. For each model we calculated Akaike’s Information Criterion (AIC) and their AIC weight (i.e. the relative likelihood) to compare between the three hypotheses. We called the inheritance of a variant as sex-linked if the relative likelihood of X- or Y-linkage exceeded 80%. We quantified the performance of this approach by calling the inheritance for variants called on the autosomes (autosomal: 98.8%, X-linked: 0.8%, Y-linked: 0. 3%), confirming that a sex-linked result is highly informative for placement on LG12. In addition, we visualized the inheritance of 100,000 variants along LG12 showing patterns of sex-linked inheritance outside the pseudo-autosomal region (Fig S32).

### Expression analysis

Following the end of selection, we sampled three replicate male and female pools of seven individuals each from each resulting post-selection population. RNA was extracted from whole tail tissue (posterior to the anal fin) using Qiagen RNeasy Mini kits (Qiagen, USA). Long-read sequencing was conducted using PacBio Revio (Pacific Biosciences, USA) and reads were subsequently processed with the PacBio IsoSeq pipeline. Reads were mapped to the female guppy reference using minimap2^97^ after clustering. We used FLAIR^98^ to further collapse redundant reads and quantify expression using counts per million (CPM). Five copies of the *texim* gene on the Y chromosome were identified by mapping Y-linked genotypes (as determined above) to collapsed full-length RNA reads, of which one (copy “1889”) was expressed at >1 CPM. Finally, we genotyped the 297 sequenced individuals for the presence of copy 1889 if they possessed the Y-linked genotype for > 50% of the SNPs in this copy.

### Software

Unless stated otherwise above, all analyses were performed in the R statistical environment^99^.

## References

1. Stephenson, J. F., Stevens, M., Troscianko, J. & Jokela, J. The Size, Symmetry, and Color Saturation of a Male Guppy’s Ornaments Forecast His Resistance to Parasites. Am. Nat. 196, 597–608 (2020).

2. Karino, K., Shinjo, S. & Sato, A. Algal-searching ability in laboratory experiments reflects orange spot coloration of the male guppy in the wild. Behaviour 144, 101–113 (2007).

3. Evans, J. P., Kelley, J. L., Bisazza, A., Finazzo, E. & Pilastro, A. Sire attractiveness influences offspring performance in guppies. Proc. R. Soc. Lond. B Biol. Sci. 271, 2035–2042 (2004).

4. Endler, J. A. Natural selection on color patterns in *Poecilia reticulata*. Evolution 34, 76–91 (1980).

5. Potter, T. et al. Female preference for rare males is maintained by indirect selection in Trinidadian guppies. Science 380, 309–312 (2023).

6. Houde, A. E. Mate choice based upon naturally occurring color-pattern variation in a guppy population. Evolution 41, 1–10 (1987).

7. Houde, A. E. & Endler, J. A. Correlated evolution of female mating preferences and male color patterns in the guppy *Poecilia reticulata*. Science 248, 1405–1408 (1990).

8. Haskins, C. P., Haskins, E. F., McLaughlin, J. J. A. & Hewitt, R. E. Polymorphism and population structure in Lebistes reticulatus, an ecological study. in Vertebrate speciation (1961).

9. Kemp, D. J., Reznick, D. N., Grether, G. F. & Endler, J. A. Predicting the direction of ornament evolution in Trinidadian guppies (*Poecilia reticulata*). Proc. R. Soc. B Biol. Sci. 276, 4335–4343 (2009).

10. Herdegen-Radwan, M., Cattelan, S., Buda, J., Raubic, J. & Radwan, J. What do orange spots reveal about male (and female) guppies? A test using correlated responses to selection. Evolution 75, 3037–3055 (2021).

11. Brooks, R. & Endler, J. A. Direct and indirect sexual selection and quantitative genetics of male traits in guppies (*Poecilia reticulata*). Evolution 55, 1002–1015 (2001).

12. Morris, J., Darolti, I., van der Bijl, W. & Mank, J. E. High-resolution characterization of male ornamentation and re-evaluation of sex linkage in guppies. Proc. R. Soc. B Biol. Sci. 287, 20201677 (2020).

13. Hughes, K. A., Houde, A. E., Price, A. C. & Rodd, F. H. Mating advantage for rare males in wild guppy populations. Nature 503, 108–110 (2013).

14. Farr, J. A. Male Rarity or Novelty, Female Choice Behavior, and Sexual Selection in the Guppy, Poecilia reticulata Peters (Pisces: Poeciliidae). Evolution 31, 162–168 (1977).

15. Parichy, D. M. & Spiewak, J. E. Origins of adult pigmentation: diversity in pigment stem cell lineages and implications for pattern evolution. Pigment Cell Melanoma Res. 28, 31–50 (2015).

16. Gordon, S. P., López-Sepulcre, A. & Reznick, D. N. Predation-associated differences in sex linkage of wild guppy coloration. Evolution 66, 912–918 (2012).

17. Toomey, M. B. et al. A mechanism for red coloration in vertebrates. Curr. Biol. 32, 4201–4214.e12 (2022).

18. Lindholm, A. K. & Breden, F. Sex Chromosomes and Sexual Selection in Poeciliid Fishes. Am. Nat. 160, (2002).

19. Tripathi, N. et al. Genetic linkage map of the guppy, Poecilia reticulata, and quantitative trait loci analysis of male size and colour variation. Proc. R. Soc. B Biol. Sci. 276, 2195–2208 (2009).

20. Nanda, I. et al. Sex chromosome polymorphism in guppies. Chromosoma 123, 373–383 (2014).

21. Kirpichnikov, V. S. Genetic Bases of Fish Selection. (Berlin : Springer, 1981).

22. Gordon, S. P., López-Sepulcre, A., Rumbo, D. & Reznick, D. N. Rapid changes in the sex linkage of male coloration in introduced guppy populations. Am. Nat. 189, 196–200 (2017).

23. Kruuk, L. E. B. Estimating genetic parameters in natural populations using the ‘animal model’. Philos. Trans. R. Soc. Lond. B. Biol. Sci. 359, 873–890 (2004).

24. Winge, Ø. One-sided masculine and sex-linked inheritance inLebistes reticulatus. J. Genet. 12, 145–162 (1922).

25. Winge, Ø. & Ditlevsen, E. Colour inheritance and sex determination in Lebistes. Heredity 1, 65–83 (1947).

26. Haskins, C. P., Young, P., Hewitt, R. E. & Haskins, E. F. Stabilised heterozygosis of supergenes mediating certain Y-linked colour patterns in populations of Lebistes Reticulatus. Heredity 25, 575–589 (1970).

27. Parichy, D. M., Ransom, D. G., Paw, B., Zon, L. I. & Johnson, S. L. An orthologue of the kit-related gene fms is required for development of neural crest-derived xanthophores and a subpopulation of adult melanocytes in the zebrafish, Danio rerio. Development 127, 3031–3044 (2000).

28. Wilkins, A. S., Wrangham, R. W. & Fitch, W. T. The “Domestication Syndrome” in Mammals: A Unified Explanation Based on Neural Crest Cell Behavior and Genetics. Genetics 197, 795–808 (2014).

29. Liley, N. R. Ethological Isolating Mechanisms in Four Sympatric Species of Poeciliid Fishes. Behav. Suppl. 1, 1–197 (1966).

30. Houde, A. E. Sex, Color, and Mate Choice in Guppies. (Princeton University Press, 1997).

31. Brooks, R. & Endler, J. A. Female Guppies Agree to Differ: Phenotypic and Genetic Variation in Mate-Choice Behavior and the Consequences for Sexual Selection. Evolution 55, 1644–1655 (2001).

32. Schroff, F., Kalenichenko, D. & Philbin, J. FaceNet: A unified embedding for face recognition and clustering. in 2015 IEEE Conference on Computer Vision and Pattern Recognition (CVPR) 815–823 (2015). doi:10.1109/CVPR.2015.7298682.

33. Almeida, P. et al. Divergence and Remarkable Diversity of the Y Chromosome in Guppies. Mol. Biol. Evol. 38, 619–633 (2021).

34. Du, K. et al. Identification of the male-specific region on the guppy Y Chromosome from a haplotype-resolved assembly. Genome Res. (2025) doi:10.1101/gr.279582.124.

35. Wright, A. E. et al. Convergent recombination suppression suggests role of sexual selection in guppy sex chromosome formation. Nat. Commun. 8, 14251 (2017).

36. Darolti, I. et al. Extreme heterogeneity in sex chromosome differentiation and dosage compensation in livebearers. Proc. Natl. Acad. Sci. 116, 19031–19036 (2019).

37. Künstner, A. et al. The Genome of the Trinidadian Guppy, Poecilia reticulata, and Variation in the Guanapo Population. PLOS ONE 11, e0169087 (2016).

38. Fraser, B. A. et al. Improved Reference Genome Uncovers Novel Sex-Linked Regions in the Guppy (Poecilia reticulata). Genome Biol. Evol. 12, 1789–1805 (2020).

39. Barriga, E. H. & Mayor, R. Chapter Nine - Embryonic Cell–Cell Adhesion: A Key Player in Collective Neural Crest Migration. in Current Topics in Developmental Biology (ed. Yap, A. S.) vol. 112 301–323 (Academic Press, 2015).

40. Yin, L. et al. Epidermal gene expression and ethnic pigmentation variations among individuals of Asian, European and African ancestry. Exp. Dermatol. 23, 731–735 (2014).

41. Yong, L., Croft, D. P., Troscianko, J., Ramnarine, I. W. & Wilson, A. J. Sensory-based quantification of male colour patterns in Trinidadian guppies reveals no support for parallel phenotypic evolution in multivariate trait space. Mol. Ecol. 31, 1337–1357 (2022).

42. Hoek, K. S. et al. Novel MITF targets identified using a two-step DNA microarray strategy. Pigment Cell Melanoma Res. 21, 665–676 (2008).

43. Zhang, Y. et al. Transcriptome analyses of betta fish (*Betta splendens*) provide novel insights into fin regeneration and color-related genes. Gene 876, 147508 (2023).

44. Yu, H.-H. & Moens, C. B. Semaphorin signaling guides cranial neural crest cell migration in zebrafish. Dev. Biol. 280, 373–385 (2005).

45. Koludrovic, D. et al. Chromatin-Remodelling Complex NURF Is Essential for Differentiation of Adult Melanocyte Stem Cells. PLOS Genet. 11, e1005555 (2015).

46. Miyadai, M. et al. Pax3 and Pax7 function in combination with Mitf to generate melanophores and xanthophores in medaka and zebrafish. 2023.06.22.546052 Preprint at 10.1101/2023.06.22.546052 (2023).

47. Bhattacharya, D., Rothstein, M., Azambuja, A. P. & Simoes-Costa, M. Control of neural crest multipotency by Wnt signaling and the Lin28/let-7 axis. eLife 7, e40556 (2018).

48. Takahashi, K. et al. A globin-family protein, Cytoglobin 1, is involved in the development of neural crest-derived tissues and organs in zebrafish. Dev. Biol. 472, 1–17 (2021).

49. Boer, E. F., Howell, E. D., Schilling, T. F., Jette, C. A. & Stewart, R. A. Fascin1-Dependent Filopodia are Required for Directional Migration of a Subset of Neural Crest Cells. PLOS Genet. 11, e1004946 (2015).

50. Lin, Y. et al. Gene duplication to the Y chromosome in Trinidadian guppies. Mol. Ecol. 31, 1853–1863 (2022).

51. Bergero, R., Gardner, J., Bader, B., Yong, L. & Charlesworth, D. Exaggerated heterochiasmy in a fish with sex-linked male coloration polymorphisms. Proc. Natl. Acad. Sci. 116, 6924–6931 (2019).

52. Tomaszkiewicz, M., Chalopin, D., Schartl, M., Galiana, D. & Volff, J.-N. A multicopy Y-chromosomal SGNH hydrolase gene expressed in the testis of the platyfish has been captured and mobilized by a Helitron transposon. BMC Genet. 15, 44 (2014).

53. Sandkam, B. A. et al. Extreme Y chromosome polymorphism corresponds to five male reproductive morphs of a freshwater fish. Nat. Ecol. Evol. 5, 939–948 (2021).

54. Lindsay, C. R. et al. P-Rex1 is required for efficient melanoblast migration and melanoma metastasis. Nat. Commun. 2, 555 (2011).

55. Endler, J. A. Natural and sexual selection on color patterns in poeciliid fishes. in Evolutionary ecology of neotropical freswater fishes 95–111 (Springer, The Netherlands, 1984).

56. Kotrschal, A. et al. Artificial Selection on Relative Brain Size in the Guppy Reveals Costs and Benefits of Evolving a Larger Brain. Curr. Biol. 23, 168–171 (2013).

57. Fong, S. et al. Rapid mosaic brain evolution under artificial selection for relative telencephalon size in the guppy (Poecilia reticulata). Sci. Adv. 7, eabj4314 (2021).

58. Kotrschal, A. et al. Rapid evolution of coordinated and collective movement in response to artificial selection. Sci. Adv. 6, eaba3148 (2020).

59. Houde, A. E. Effect of artificial selection on male colour patterns on mating preference of female guppies. Proc. R. Soc. Lond. B Biol. Sci. 256, 125–130 (1994).

60. Cole, G. L. & Endler, J. A. Change in male coloration associated with artificial selection on foraging colour preference. J. Evol. Biol. 31, 1227–1238 (2018).

61. Yamamoto, T. The Medaka, Oryzias latipes, and the Guppy, Lebistes reticularis. in Handbook of Genetics: Volume 4 Vertebrates of Genetic Interest (ed. King, R. C.) 133–149 (Springer US, Boston, MA, 1975). doi:10.1007/978-1-4613-4470-4_7.

62. Houde, A. E. Sex-linked heritability of a sexually selected character in a natural population of *Poecilia reticulata* (Pisces: Poeciliidae) (guppies). Heredity 69, 229–235 (1992).

63. Inaba, M., Yamanaka, H. & Kondo, S. Pigment Pattern Formation by Contact-Dependent Depolarization. Science 335, 677–677 (2012).

64. Kottler, V. A., Fadeev, A., Weigel, D. & Dreyer, C. Pigment Pattern Formation in the Guppy, Poecilia reticulata, Involves the Kita and Csf1ra Receptor Tyrosine Kinases. Genetics 194, 631–646 (2013).

65. Kottler, V. A. et al. Adenylate cyclase 5 is required for melanophore and male pattern development in the guppy (Poecilia reticulata). Pigment Cell Melanoma Res. 28, 545–558 (2015).

66. Brush, A. H. Metabolism of carotenoid pigments in birds. FASEB J. 4, 2969–2977 (1990).

67. Hill, G. E. & Johnson, J. D. The Vitamin A–Redox Hypothesis: A Biochemical Basis for Honest Signaling via Carotenoid Pigmentation. Am. Nat. 180, E127–E150 (2012).

68. Grether, G. F., Hudon, J. & Millie, D. F. Carotenoid limitation of sexual coloration along an environmental gradient in guppies. Proc. R. Soc. Lond. B Biol. Sci. 266, 1317–1322 (1999).

69. Lopes, R. J. et al. Genetic Basis for Red Coloration in Birds. Curr. Biol. 26, 1427–1434 (2016).

70. Mundy, N. I. et al. Red Carotenoid Coloration in the Zebra Finch Is Controlled by a Cytochrome P450 Gene Cluster. Curr. Biol. 26, 1435–1440 (2016).

71. McLean, C. A. et al. Red carotenoids and associated gene expression explain colour variation in frillneck lizards. Proc. R. Soc. B Biol. Sci. 286, 20191172 (2019).

72. Pérez-Gálvez, A. & Mínguez-Mosquera, M. I. Esterification of xanthophylls and its effect on chemical behavior and bioavailability of carotenoids in the human. Nutr. Res. 25, 631–640 (2005).

73. Breithaupt, D. E., Bamedi, A. & Wirt, U. Carotenol fatty acid esters: easy substrates for digestive enzymes? Comp. Biochem. Physiol. B Biochem. Mol. Biol. 132, 721–728 (2002).

74. Paris, J. R. et al. A large and diverse autosomal haplotype is associated with sex-linked colour polymorphism in the guppy. Nat. Commun. 13, 1233 (2022).

75. Bruders, R. et al. A copy number variant is associated with a spectrum of pigmentation patterns in the rock pigeon (Columba livia). PLOS Genet. 16, e1008274 (2020).

76. Weich, K. et al. Pigment Intensity in Dogs is Associated with a Copy Number Variant Upstream of KITLG. Genes 11, 75 (2020).

77. Kokko, H., Jennions, M. D. & Houde, A. Evolution of frequency-dependent mate choice: keeping up with fashion trends. Proc. R. Soc. B Biol. Sci. 274, 1317–1324 (2007).

78. Bürkner, P.-C. Advanced Bayesian Multilevel Modeling with the R Package brms. R J. 10, 395–411 (2018).

79. Bürkner, P.-C. brms: An R Package for Bayesian Multilevel Models Using Stan. J. Stat. Softw. 80, 1–28 (2017).

80. Lenth, R. V. Emmeans: Estimated Marginal Means, Aka Least-Squares Means. (2023).

81. Bates, D., Mächler, M., Bolker, B. & Walker, S. Fitting Linear Mixed-Effects Models Using lme4. J. Stat. Softw. 67, 1–48 (2015).

82. Pérez-Escudero, A., Vicente-Page, J., Hinz, R. C., Arganda, S. & de Polavieja, G. G. idTracker: tracking individuals in a group by automatic identification of unmarked animals. Nat. Methods 11, 743–748 (2014).

83. Brooks, M. E. et al. glmmTMB Balances Speed and Flexibility Among Packages for Zero-inflated Generalized Linear Mixed Modeling. R J. 9, 378–400 (2017).

84. Carpenter, B. et al. Stan: A Probabilistic Programming Language. J. Stat. Softw. 76, (2017).

85. Wolak, M. E. nadiv: an R package to create relatedness matrices for estimating non-additive genetic variances in animal models. Methods Ecol. Evol. 3, 792–796 (2012).

86. Giovanny, C.-P. Genome assisted prediction of quantitative traits using the R package sommer. PLoS ONE 11, 1–15 (2016).

87. McInnes, L., Healy, J. & Melville, J. UMAP: Uniform Manifold Approximation and Projection for Dimension Reduction. Preprint at 10.48550/arXiv.1802.03426 (2020).

88. Melville, J. Uwot: The Uniform Manifold Approximation and Projection (UMAP) Method for Dimensionality Reduction. (2023).

89. Zhang, Z. et al. Mixed linear model approach adapted for genome-wide association studies. Nat. Genet. 42, 355–360 (2010).

90. Zhou, X. & Stephens, M. Genome-wide efficient mixed-model analysis for association studies. Nat. Genet. 44, 821–824 (2012).

91. Legarra, A., Aguilar, I. & Misztal, I. A relationship matrix including full pedigree and genomic information. J. Dairy Sci. 92, 4656–4663 (2009).

92. Li, X. & Zhu, X. Cross-Phenotype Association Analysis Using Summary Statistics from GWAS. in Statistical Human Genetics: Methods and Protocols (ed. Elston, R. C.) 455–467 (Springer, New York, NY, 2017). doi:10.1007/978-1-4939-7274-6_22.

93. Lin, Y., Darolti, I., Bijl, W. van der, Morris, J. & Mank, J. E. Extensive variation in germline de novo mutations in Poecilia reticulata. Genome Res. 33, 1317–1324 (2023).

94. Benjamini, Y. & Hochberg, Y. Controlling the False Discovery Rate: A Practical and Powerful Approach to Multiple Testing. J. R. Stat. Soc. Ser. B Methodol. 57, 289–300 (1995).

95. Krassowski, M. ComplexUpset. (2020) doi:10.5281/zenodo.3700590.

96. Baglama, J., Reichel, L. & Lewis, B. W. Irlba: Fast Truncated Singular Value Decomposition and Principal Components Analysis for Large Dense and Sparse Matrices. (2022).

97. Li, H. Minimap2: pairwise alignment for nucleotide sequences. Bioinformatics 34, 3094–3100 (2018).

98. Tang, A. D. et al. Full-length transcript characterization of SF3B1 mutation in chronic lymphocytic leukemia reveals downregulation of retained introns. Nat. Commun. 11, 1438 (2020).

99. R Core Team. R: A Language and Environment for Statistical Computing. (R Foundation for Statistical Computing, Vienna, Austria, 2023).

