## Supplemental information for "Deep learning reveals the complex genetic architecture of a highly polymorphic sexual trait"

### Supplemental methods

#### *Color pattern extraction*

We used a custom pipeline to analyze images, consisting of five steps:

Step 1: We first performed image segmentation to separate the fish from the background. We excluded the gonopodium and dorsal fin from the extracted fish; since their size is not consistent on camera, they are difficult to segment due to their transparency, and they do not exhibit ornamentation in our population. We used a convolutional neural net with a Unet[73] architecture, implemented in the *unet* R package[74], with six layers in the encoder, 10 filters in the first convolution, a dropout of 0.25 and with binary cross-entropy as the loss function. Using the segmentation mask, we then cropped and rotated the fish to an image of dimension [800 x 300 x 4] (three color channels and the segmentation mask).

Step 2: Next, we further segmented the fish image with two additional Unet models, to extract the carotenoid and melanic coloration respectively. Both used four layers in the encoder, 12 filters in the first convolution, a dropout of 0.5 and a Dice loss function.

Step 3: To be able to directly compare regions of the guppy body, we performed image alignment. We trained a convolutional neural net to place four landmarks on the fish: in the center of the eye, on the tip of the snout, and on the dorsal and ventral extremes of the boundary between the caudal peduncle and the tail fin. Instead of predicting the landmark coordinates directly, which worked poorly, the landmarks were represented as a stack of four images, each having a point (with Gaussian blur) at one landmark location. The model was then trained to predict these landmark images, taking the extracted fish from Step 1 as input. The model first has three convolutional layers, followed by three deconvolutional layers, to transform the three color channels into four heatmaps of landmark probability (Table S1). We then selected the highest activation on each output channel as the predicted landmark coordinates.

Step 4: Treating the four landmarks from Step 3 as fixed, we added sliding semi-landmarks algorithmically. We first mirrored all images of the left side of the fish, then found all pixels on the boundary of the segmentation mask from Step 1, which is the outline of the fish. These pixels were ordered into a continuous curve by treating the connection between points as a Travelling Salesman Problem. Using the fixed landmarks from Step 3, we split this curve into three: a) the outline from the tip of the snout, along the back, to the dorsal attachment of the caudal fin, b) the outline of the caudal fin, dorsal to ventral, and c) from the ventral attachment of the caudal fin to the tip of the snout. Along each of these curves we placed 50 evenly spaced semi-landmarks.

Step 5: Using the fixed landmarks from Step 3 and sliding landmarks from Step 4, we used Generalized Procrustes analysis to generate a consensus shape using package *geomorph*[75]. Using thin-plate spline transformation, we then warped our fish images to the consensus shape with package *Morpho*[76]. This warping is applied to the segmented fish from Step 1, and the segmented ornamentation from Step 2. Since these are continuous transformations, we then rasterized the warped output by taking color averages along a grid, resulting in images of dimension [500 x 140 x 4].

By comparing the number of pixels in the extracted fish mask from Step 1, and the number of pixels in the extracted carotenoid and melanin masks from Step 2 (i.e. before warping), we calculated the amount of coloration as a percentage of the body area. For use in our selection paradigm, we calculated the mean across multiple photos of each side of each fish, and then the mean across the sides.

For manual segmentation for Step 1, we annotated the fish's outline with splines using the *StereoMorph* package[77]. For manual color segmentation in Step 2, we used the free and fuzzy select tools in the GNU Image Manipulation Program (GIMP). For black segmentation, we only selected distinct color patches of concentrated pigmentation, which are always present. We excluded the various facultatively expressed darker areas, sometimes referred to as "fuzzy black"[11,30].

#### *Color calibration*

Fish were photographed while swimming in a photarium. For the P and $F_1$ generations, we used a glass photarium measuring 16 x 85 x 100 mm (internal length x width x height), with an approximate water level of 3 cm. For generations $F_2$ and $F_3$, we instead used a thin plastic 54 mm cuboid box with an internal sliding white wall. In this photarium (following (*74*)) the fish can be restricted to a very narrow area in the front by sliding the white wall forward. Since the change in photarium led to small changes in color characteristics, we performed color calibration to match the two setups. To this end, we photographed 15 fish with both methods, and extracted the fish from the images. We then generated look-up-tables (LUTs) by matching color quantiles from these images in Lab color space. Photographs from the new photorarium were then adjusted to match the old color profile using the LUTs prior to analysis.

*Effect size simulation*

To evaluate whether the response to selection was unexpected in magnitude, we compared the observed differences in orange ornament incidence with predictions from a modified breeder's equation. The modified breeder's equation separates autosomal, X- and Y-linked heritability, models selection on males only, and is parametrized using the difference in population means. These changes result in the following equation, for the difference in population means $\Delta\bar{X}$ in generation $t+1$:

$$\Delta\bar{X}_{t+1} = \left(\frac{1}{2}h_A^2 + h_Y^2\right)\left(S_t^{\downarrow} - S_t^{\uparrow}\right) + \frac{1}{3}h_X^2\left(S_{t-1}^{\downarrow} - S_{t-1}^{\uparrow}\right) + \Delta\bar{X}_t$$

Where $h_A^2$, $h_X^2$ and $h_Y^2$ are the autosomal, X- and Y-linked heritabilities, and $S_t^{\downarrow}$ and $S_t^{\uparrow}$ are the selection differentials in generation $t$ for the down- and up-selected lines, respectively. As we are predicting ornament incidences, all parameters are on the link (liability) scale. We compared predictions and observations (Figure S6) for each replicate separately, removing generations where the incidence was 0 or 1. Selection differentials before the onset of artificial selection are assumed to be 0. We evaluated accuracy using the squared Pearson correlation coefficient ($r^2$ = 0.83).

*Reference genome comparison*

To evaluate the potential for reference bias, we compared mapping rates between female (*36*) and male (*37*) reference genomes. Reference bias, where reads from some individuals map better to the reference than reads from other individuals, should result in increased variance in mapping rates between individuals. We therefore compared the variance in mapping rates between these two genome assemblies.

First, we compared mapping rates of all reads to the chromosome-level scaffolds of these two reference genomes, after filtering on mapping quality to match our genomic analyses. As expected from mapping bias of Y haplotypes to the male reference, variance in mapping rates to the male reference was 86% higher (F-test, F = 1.86, df = 294, p < 0.001). We also found that mapping rates to the female reference were higher for 97% of individuals, and higher on average (paired t-test, t = 24.501, df = 294, p-value < 0.001).

Supplemental figures

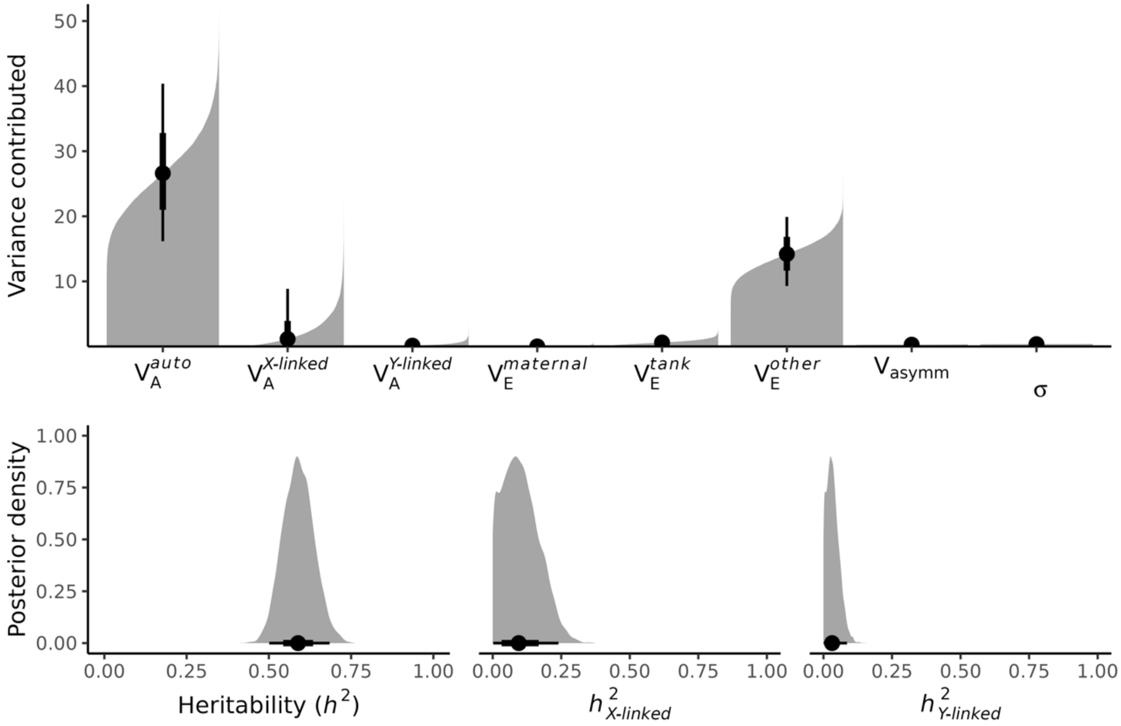


**Figure S1: Variance components of total amount of orange color**, as estimated by a Bayesian animal model. Top panel shows cumulative posterior density bar graphs, bottom panel shows posterior distributions. Posterior summaries are shown as posterior means (point) and 66% (thick lines) and 95% (thin lines) credible intervals.

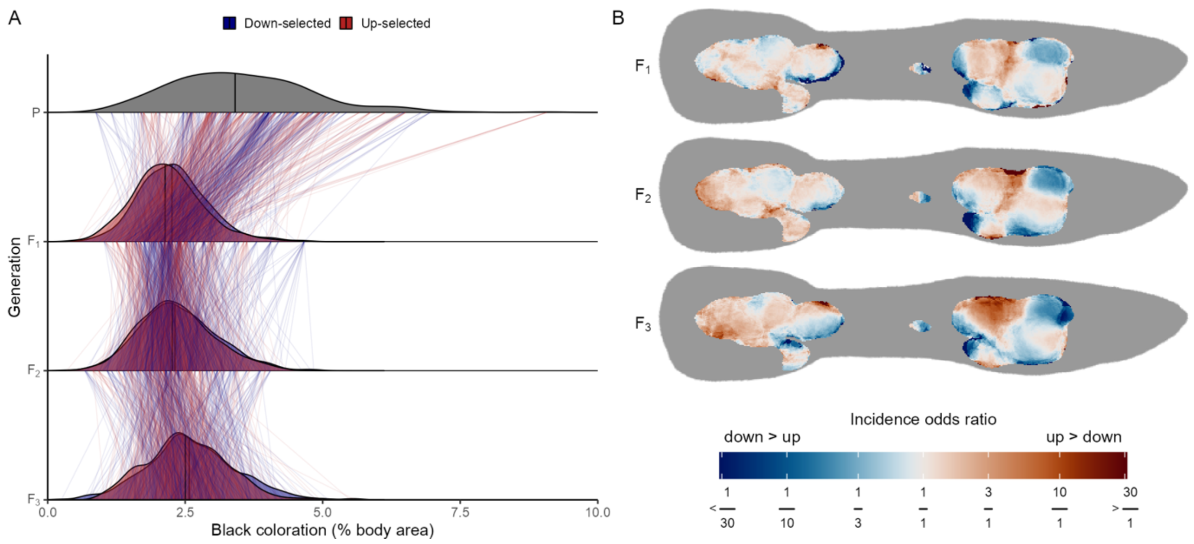


**Figure S2: Correlated effects of selection on orange area on the incidence of black coloration.** A) Density plots show the distribution of the percent of black coloration per generation and selection regime. Thin lines between generations connect fathers and sons. Thick lines inside the densities show the median value of color area. B) Heatmaps illustrating the effect of selection across the body. Each heatmap cell is colored by the log odds ratio (as estimated by a generalized linear mixed model), illustrating the relative odds that a male has black color at that location. Body positions where the incidence of black color is less than 1% are colored grey.

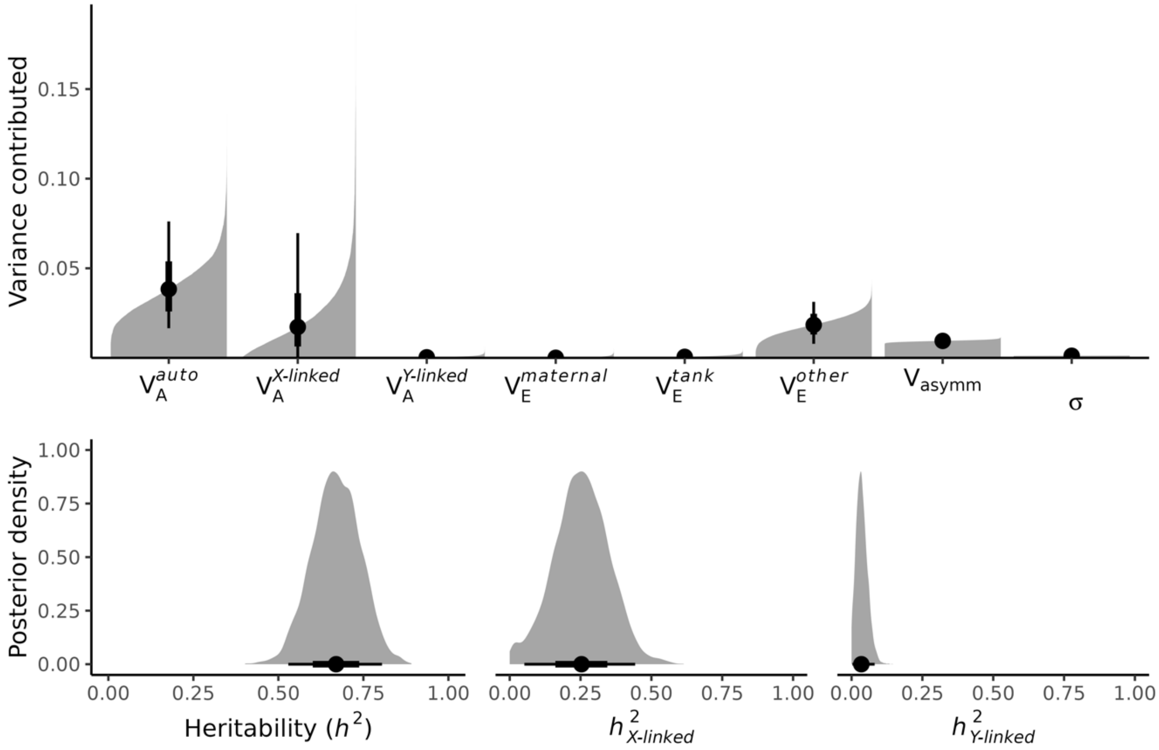


**Figure S3: Variance components of total amount of black color**, as estimated by a Bayesian animal model. Top panel shows cumulative posterior density bar graphs, bottom panel shows posterior distributions. Posterior summaries are shown as posterior means (point) and 66% (thick lines) and 95% (thin lines) credible intervals.

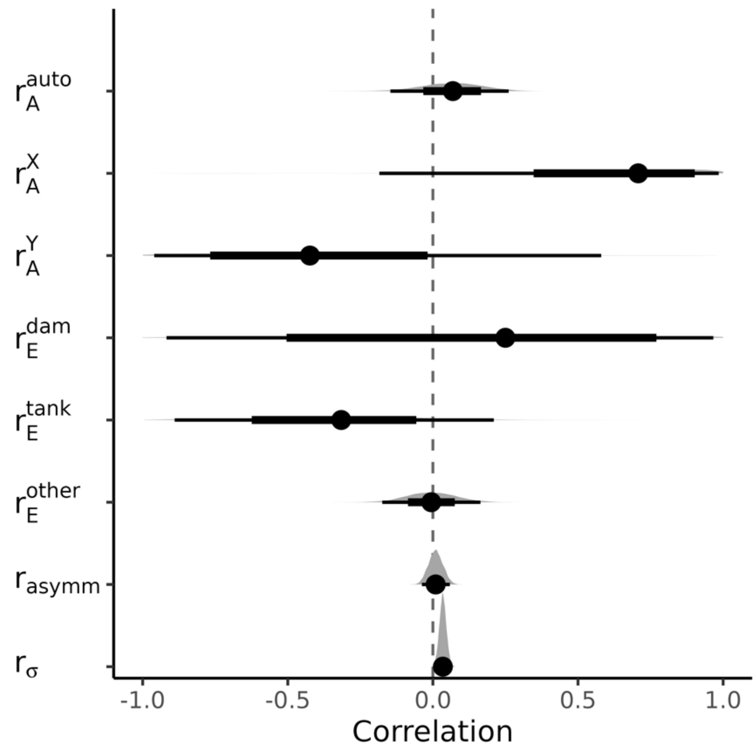


**Figure S4: Posterior distributions of the correlation of orange and black variance components** (Figs S1&S2). Posterior summaries are shown as posterior means (point) and 66% (thick lines) and 95% (thin lines) credible intervals.

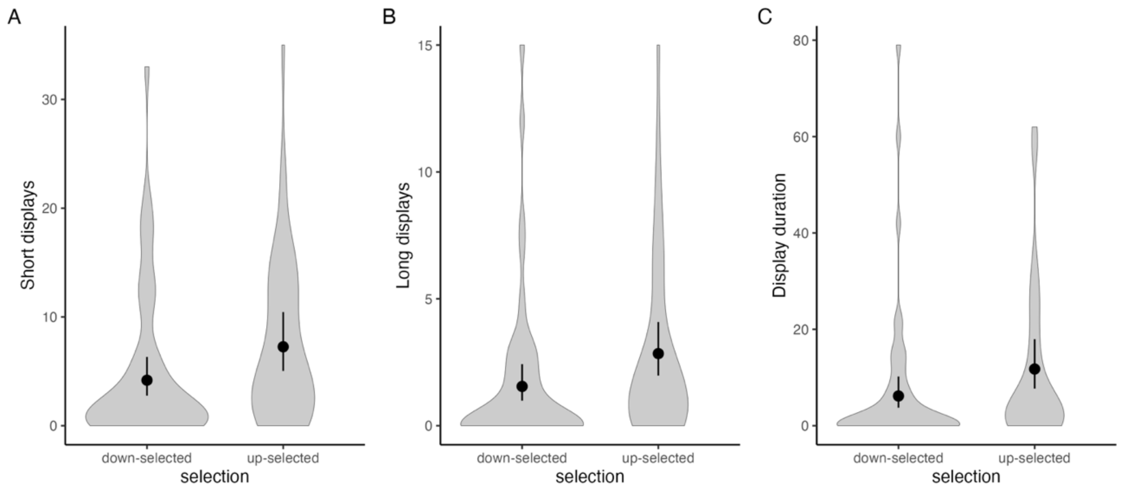


**Fig S5:** The effect of artificial selection for orange coloration on display behavior of male guppies. Comparison of a) frequency of short displays; b) frequency of long displays; and c) duration of display in down-selected (n = 46) and up-selected (n = 49) males from the $F_3$ generation. Violin plots show the kernel densities of observations, and point estimates and error bars show the point estimate and 95% confidence intervals for the groups means estimated by a linear mixed model.

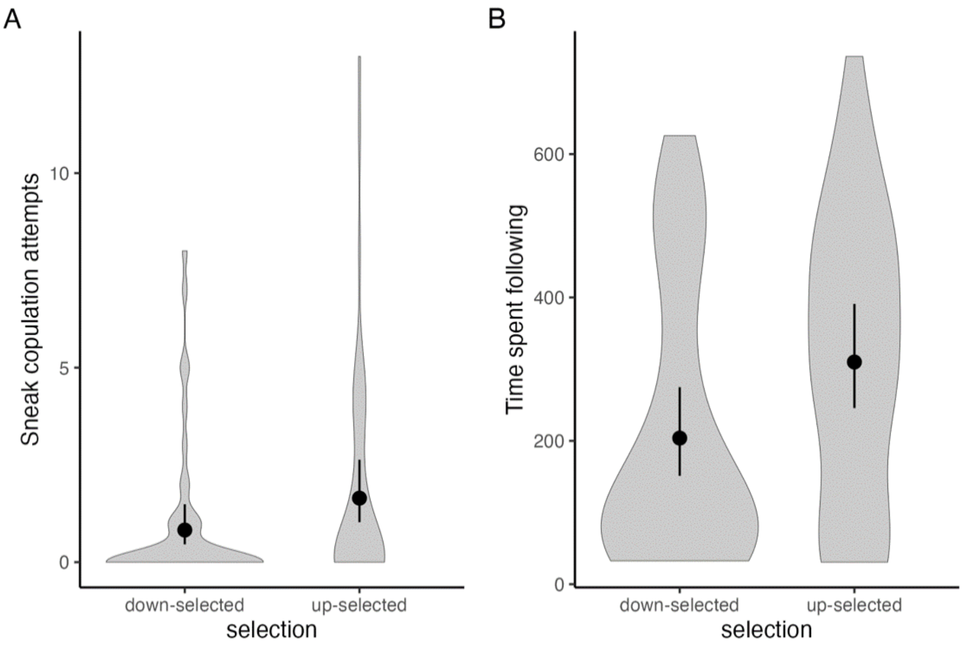


**Fig S6:** The effect of artificial selection for orange coloration on coercive sexual behavior of male guppies. Comparison of a) sneak copulation attempts and b) time spent following the female in of down-selected (n = 46) and up-selected (n = 49) males from the $F_3$ generation. Violin plots show the kernel densities of observations, and point estimates and error bars show the point estimate and 95% confidence intervals for groups mean as estimated by a linear mixed model.

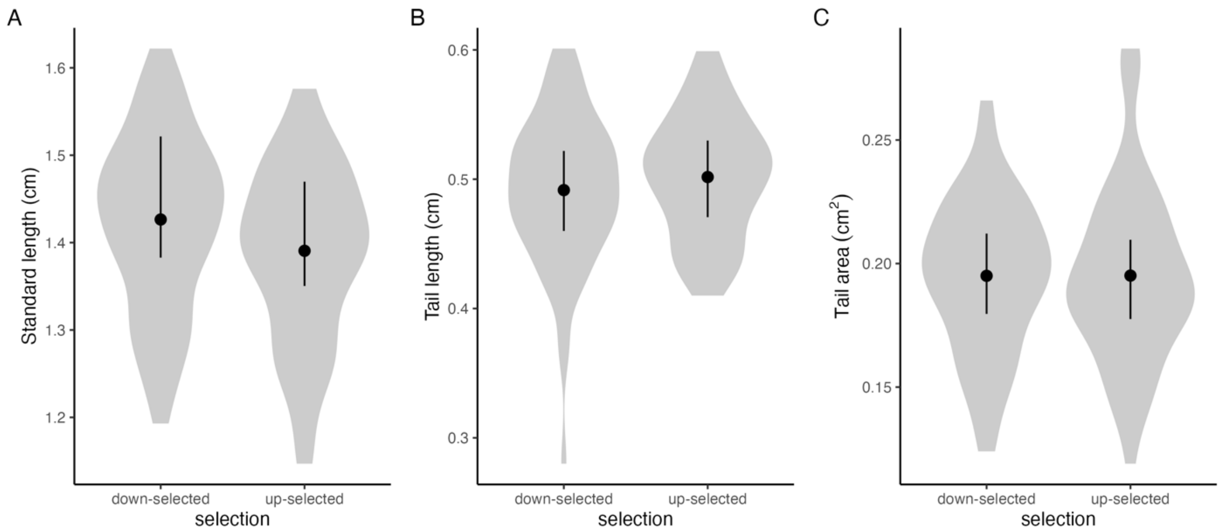


**Fig S7:** The effect of artificial selection for orange coloration on gross morphology of male guppies. Comparison of a) standard length b) tail length and c) tail area in male guppies following three generations of artificial selection for down-selected (n = 63) and up-selected (n = 62) males. Violin plots show the kernel densities of observations while point estimates and error bars show the posterior median and 95% credible intervals for the groups means estimated by a linear mixed model.

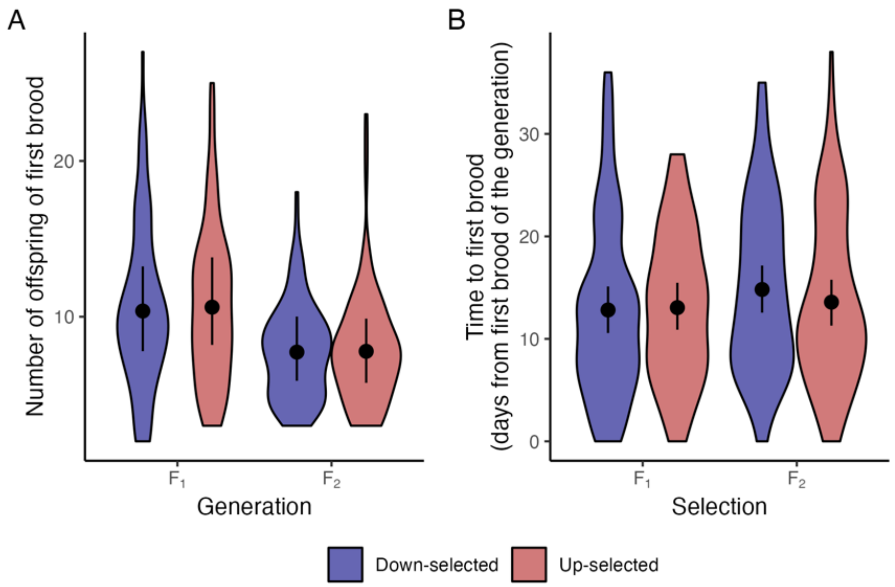


**Fig S8**: The effect of selection on orange coloration on life history parameters. A) shows the fecundity of females, as the number of offspring in their first brood. B) shows the time between pairing and the first brood (day 0 is the first brood of the generation). Violin plots show the kernel densities of observations while point estimates and error bars show the posterior median and 95% credible intervals for the groups means estimated by a linear mixed model.

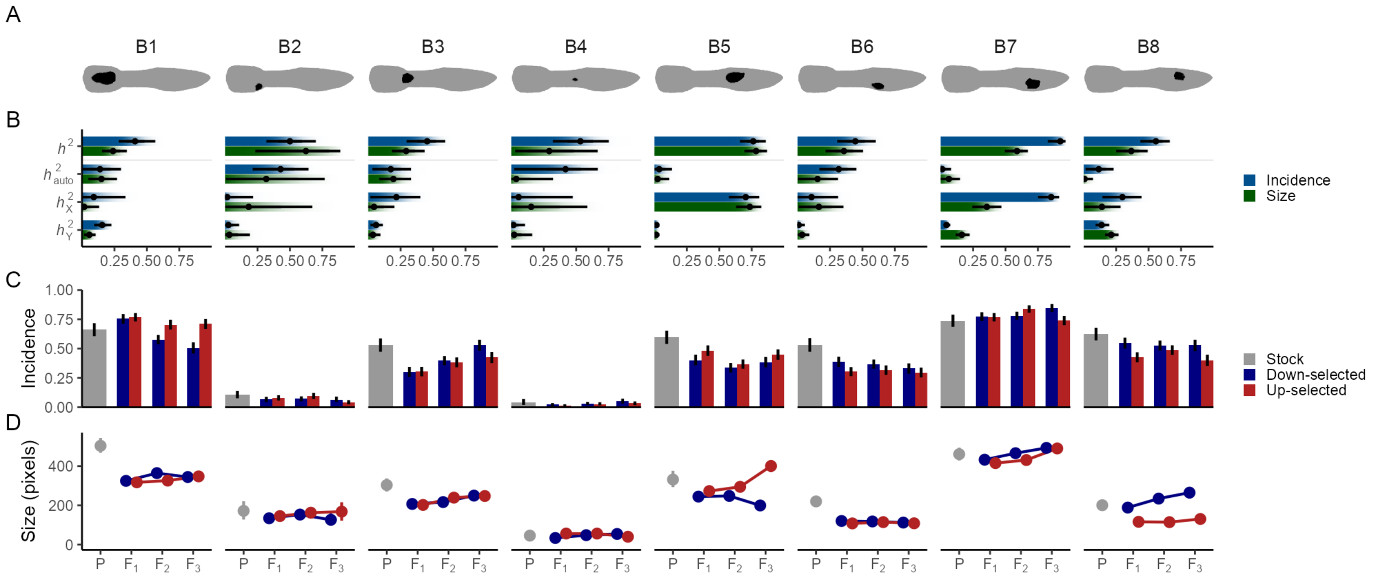


**Fig S9: Black ornaments are heritable but responded only weakly to selection on orange color**. A) Pictograms of eight black ornaments. B) Heritabilities of the incidence and size (when present) of each ornament. Dots and lines reflect point estimates and 95% credible intervals, and the gradient bars show the cumulative posterior density. C&D) Effect of selection on the incidence and size of black ornaments. Error bars show 95% bootstrapped confidence intervals, and points represent means. X-axes show consecutive generations.

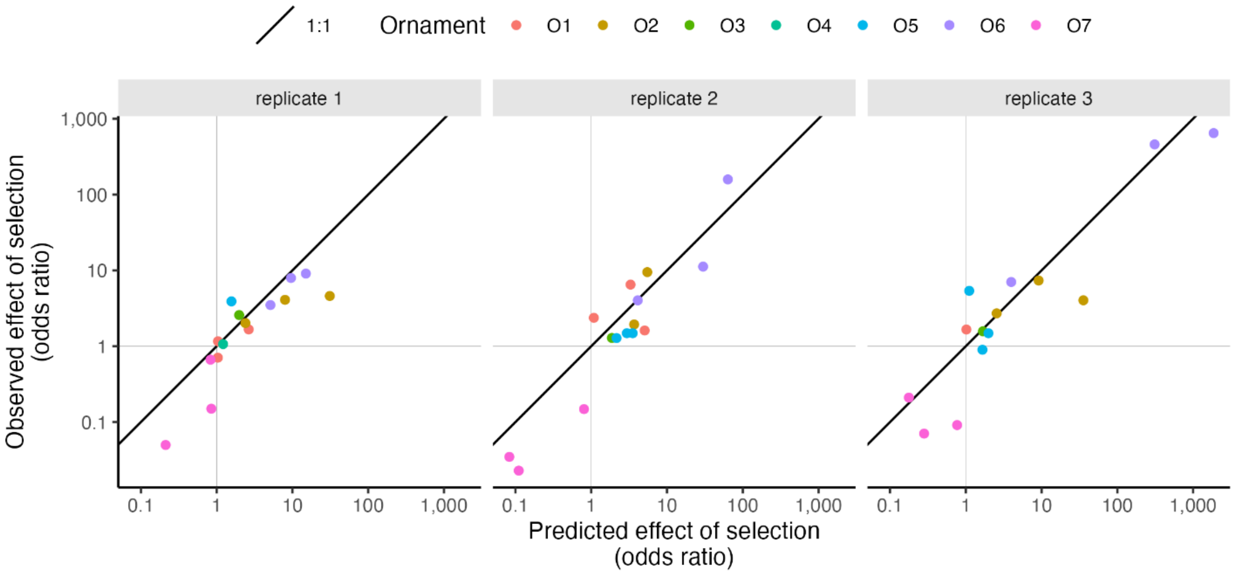


**Figure S10: Observed effect sizes of artificial selection on the incidence of orange ornaments follow expectations arising from the breeder's equation** ($r^2$ = 0.83). Points are colored by ornament (see Fig 4). The solid line is the 1:1 relationship, where the thin horizontal and vertical lines indicate no effect of selection (odds ratio = 1).

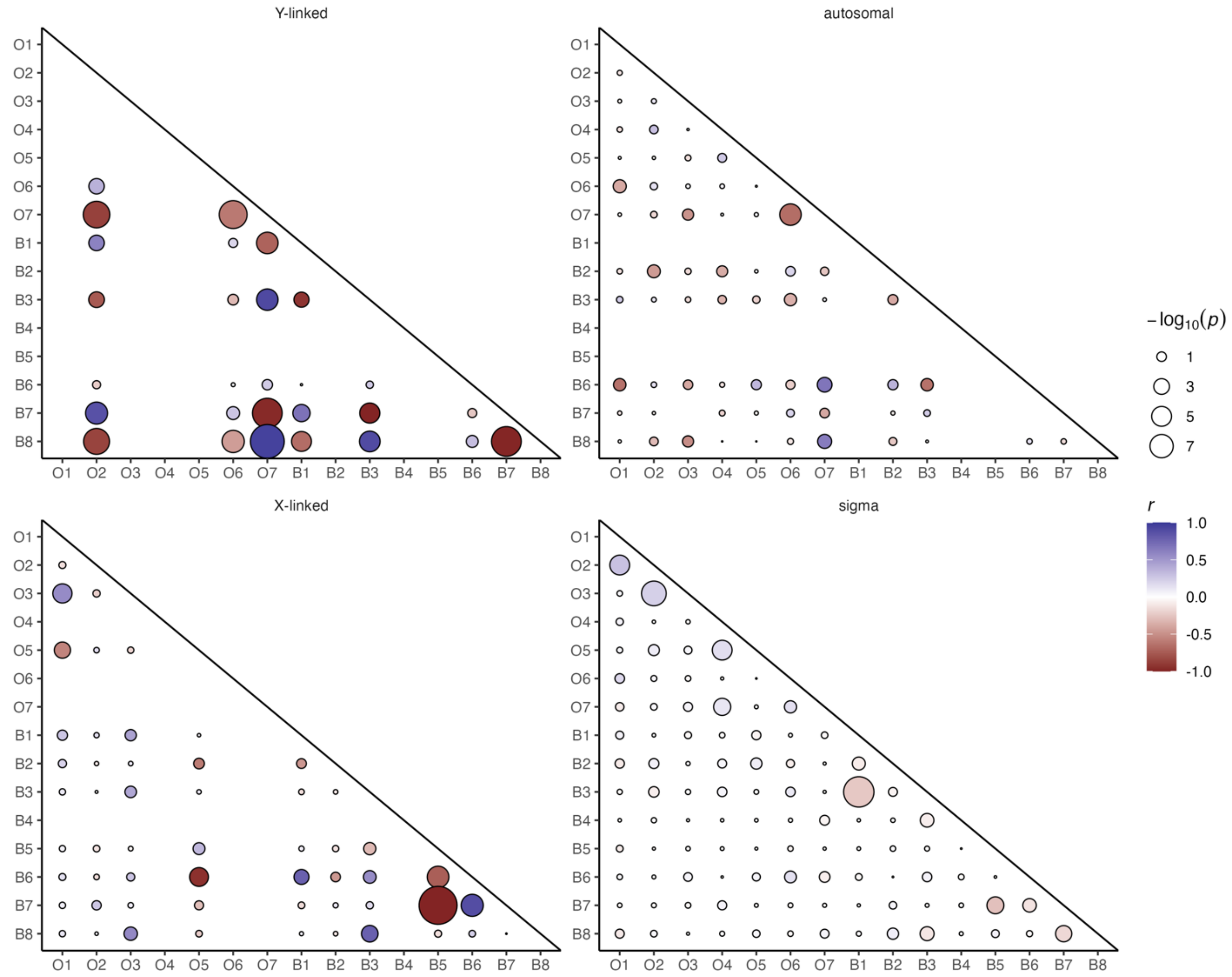


**Fig S11: Pairwise correlations of the presence of ornaments**, separated in the Y-linked, X-linked and autosomal components, and the environment (sigma). The asymptotic -log10(p-value) of the correlation is shown as the point size, with the color reflecting the estimating correlation coefficient *r*. Correlations are only shown if the variance component was significantly greater than zero for both ornaments.

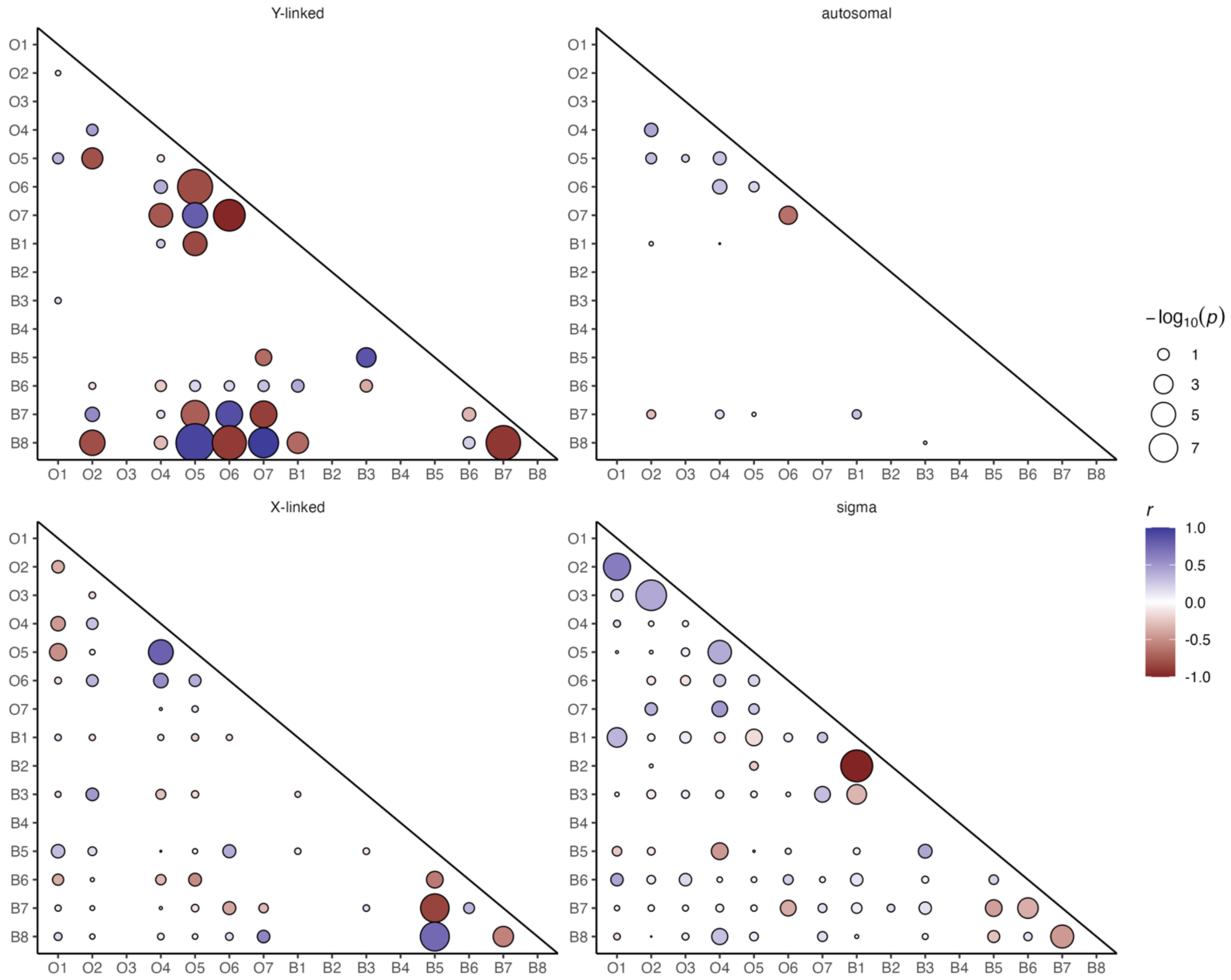


**Fig S12: Pairwise correlations of the size of ornaments**, separated in the Y-linked, X-linked and autosomal components, and the environment (sigma). The asymptotic -log10(p-value) of the correlation is shown as the point size, with the color reflecting the estimating correlation coefficient *r*. Correlations are only shown if the variance component was significantly greater than zero for both ornaments. Models only included individuals with both ornaments present.

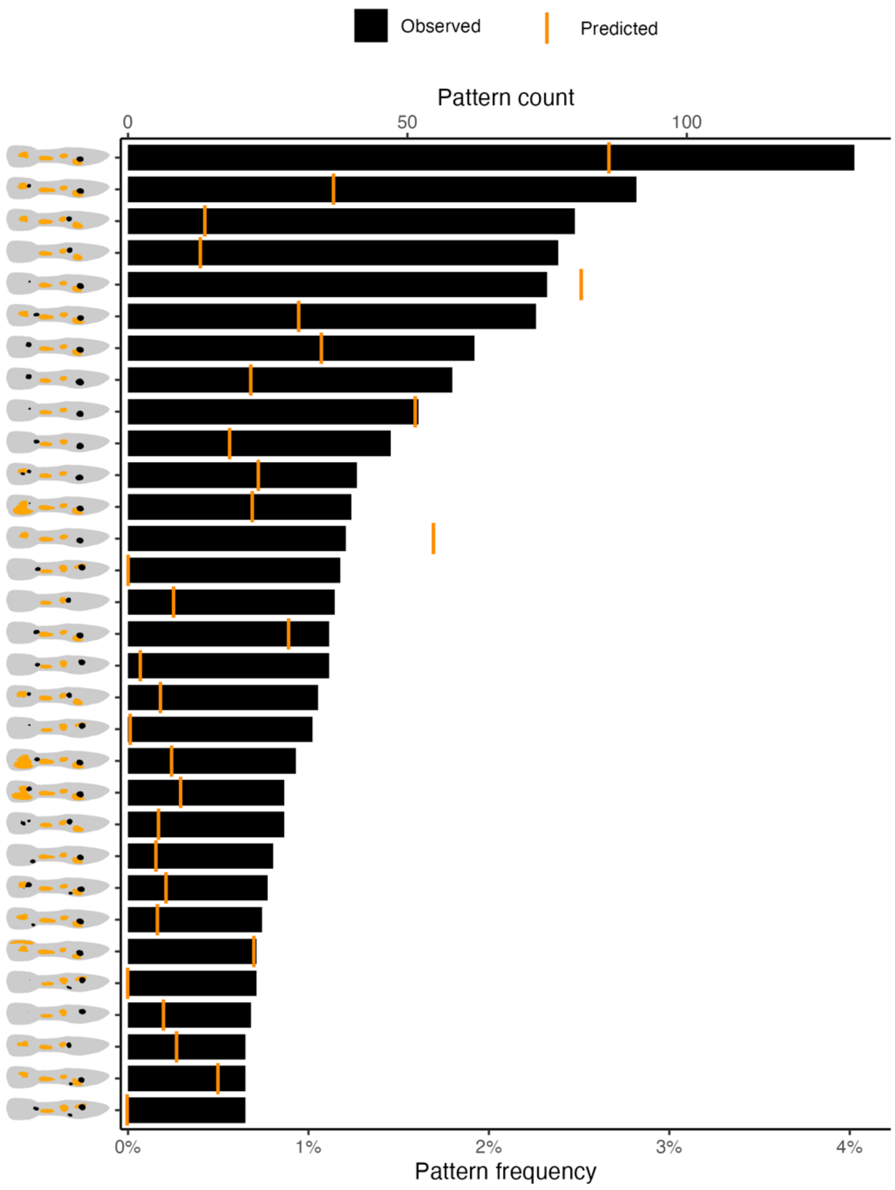


**Fig S13**: Top 30 most common ornament combinations. The y-axis shows the median color pattern for each combination of ornaments, while x-axis shows its count (top) and frequency in the population (bottom). The bars indicate the observed frequency, while the orange lines show the frequency predicted from the incidence of each of the pattern's ornaments (see Fig S14).

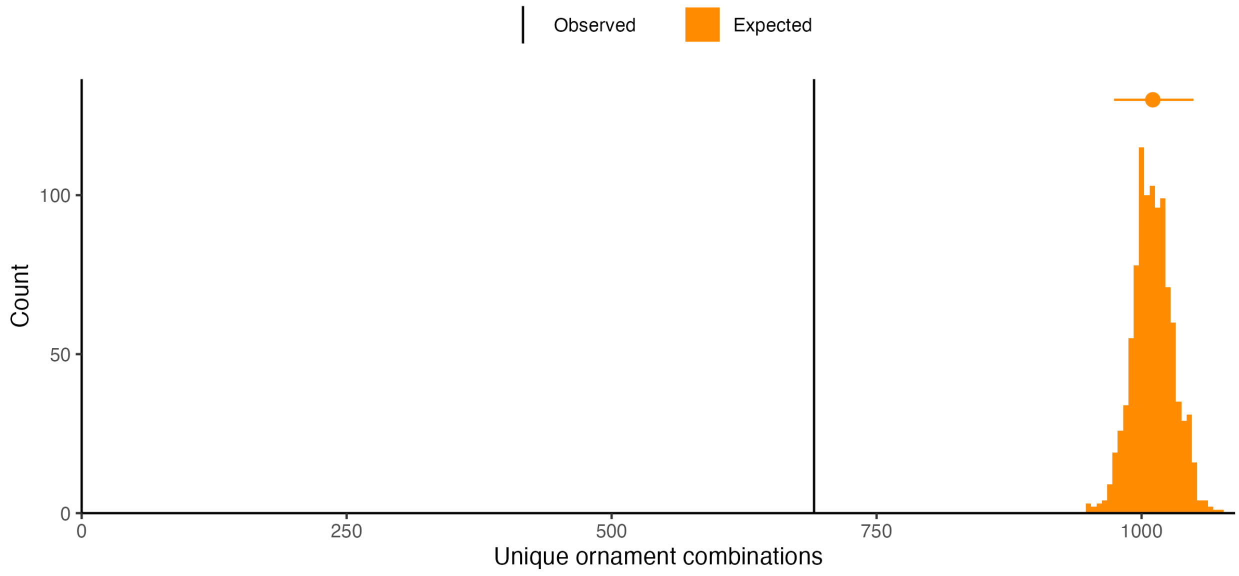


**Fig S14**: The number of unique ornament combinations in the full dataset of all males (black line), compared the expectation derived from ornament incidence (orange). The expected distribution was calculated by randomly generating 3,229 males and their ornaments with P(incidence), repeated 1,000 times. The point and errorbar denote the mean and 95% confidence interval.

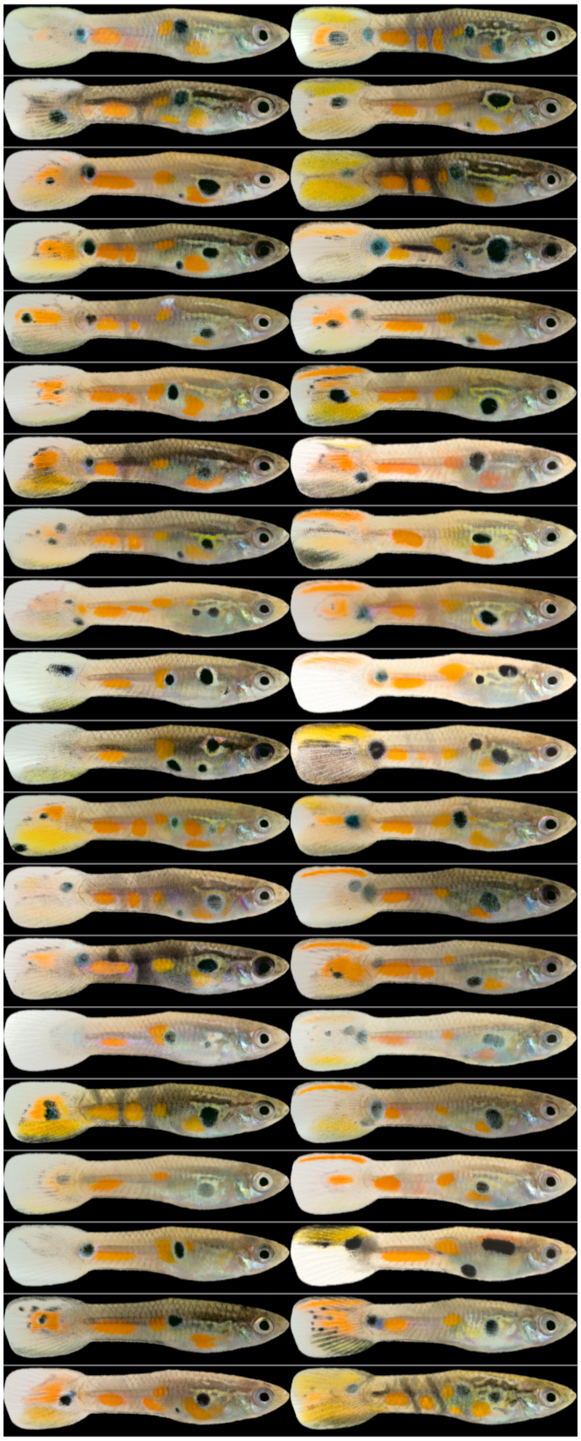

**Fig S15: Ornament diversity in the tail fin.** Displayed are randomly selected examples of fish with an absent (left) or present (right) ornament O1 (top of the tail fin). Note that the fish with a present ornament O1 have variety of tail ornamentation, possibly indicating spatially overlapping ornaments.

A

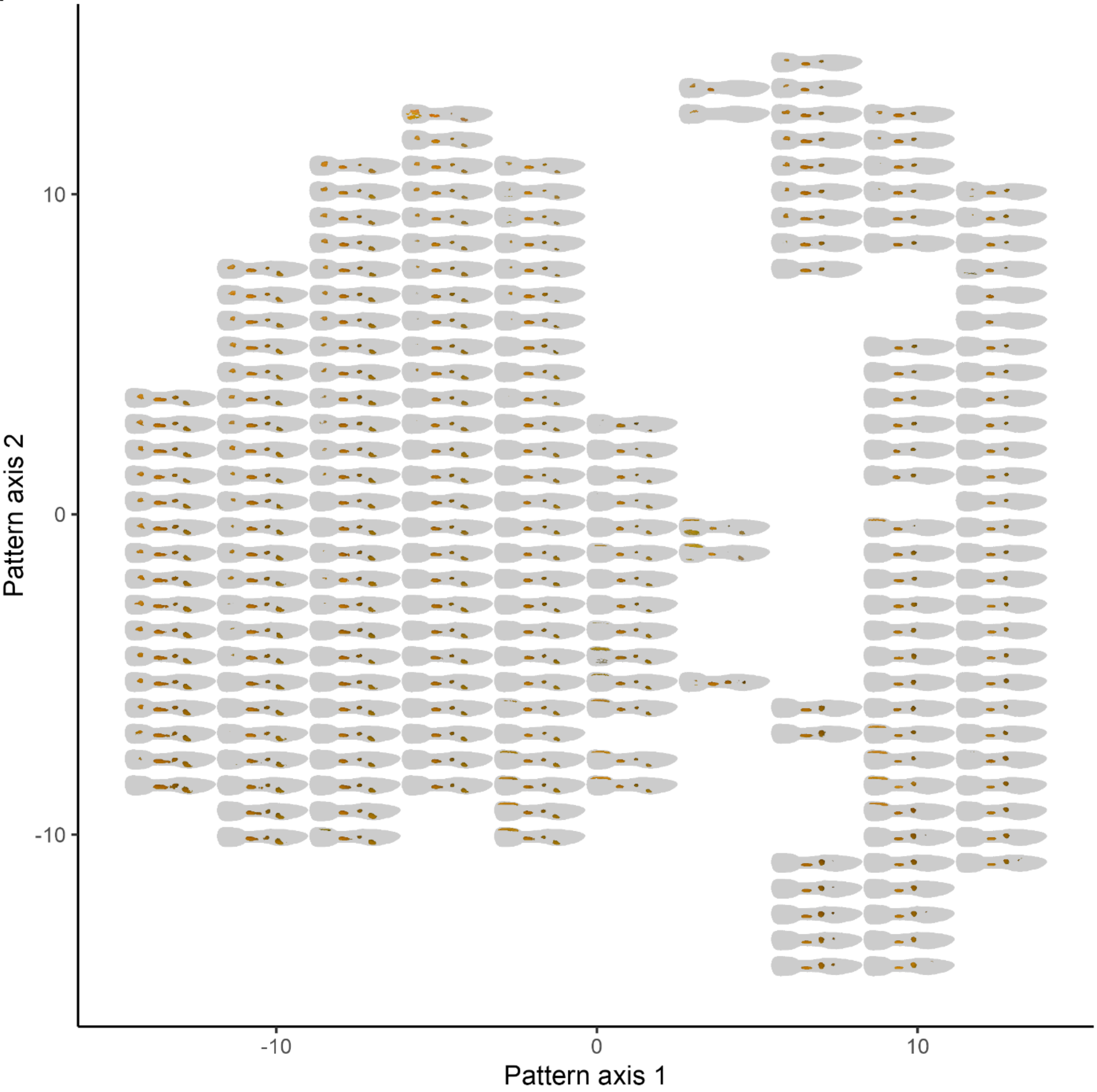


(continues on next page…)

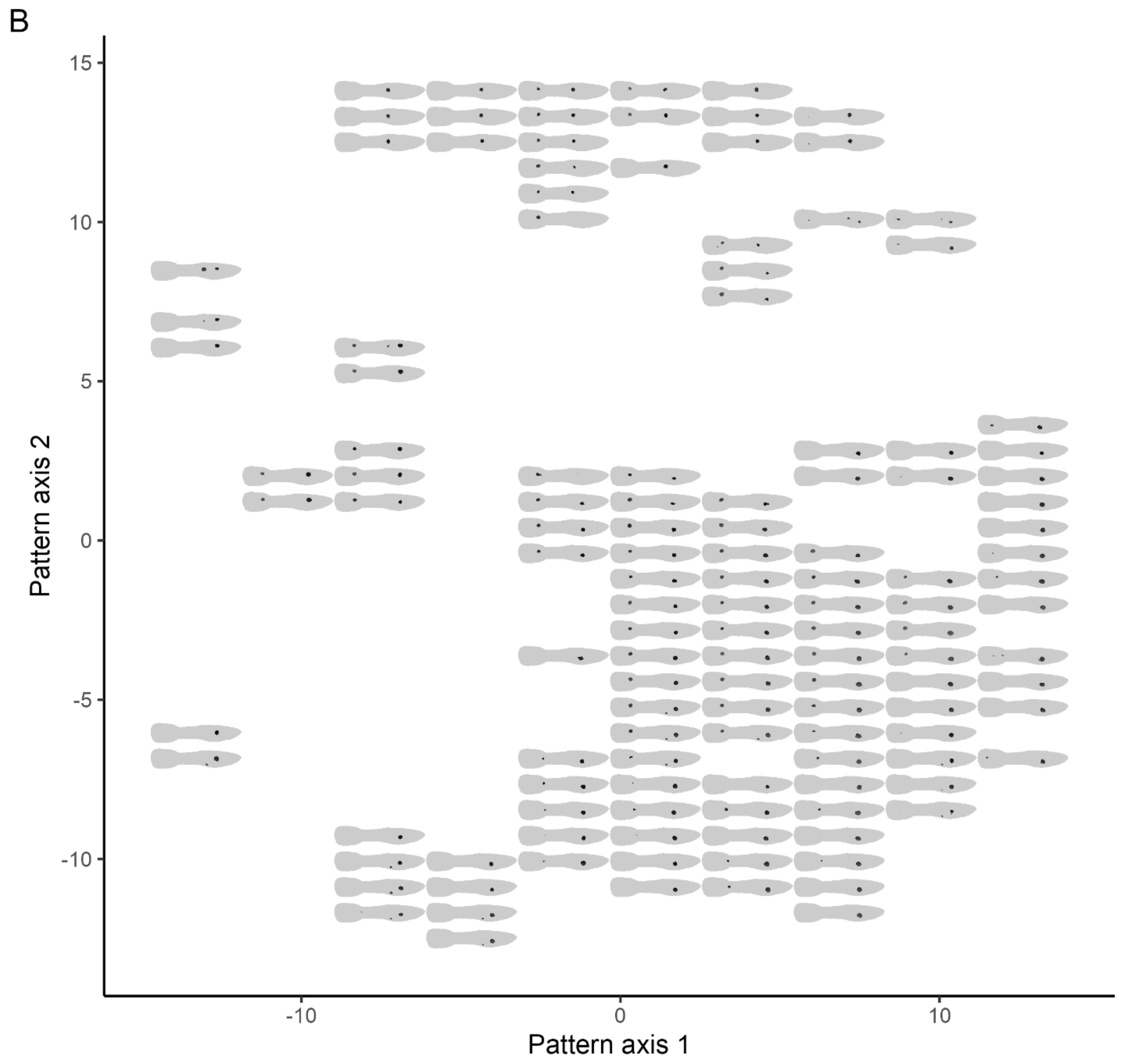


**Figure S16: Patternspaces provide holistic summaries of pattern variation.** A) Axes are the UMAP reduced representation of five-dimensional orange patternspace. Fish pictograms depict the median orange color pattern for all images falling within the grid position of the picogram. Empty locations have no images in that part of patternspace (see Fig S11 for image distribution). B) As in A, but for black patternspace.

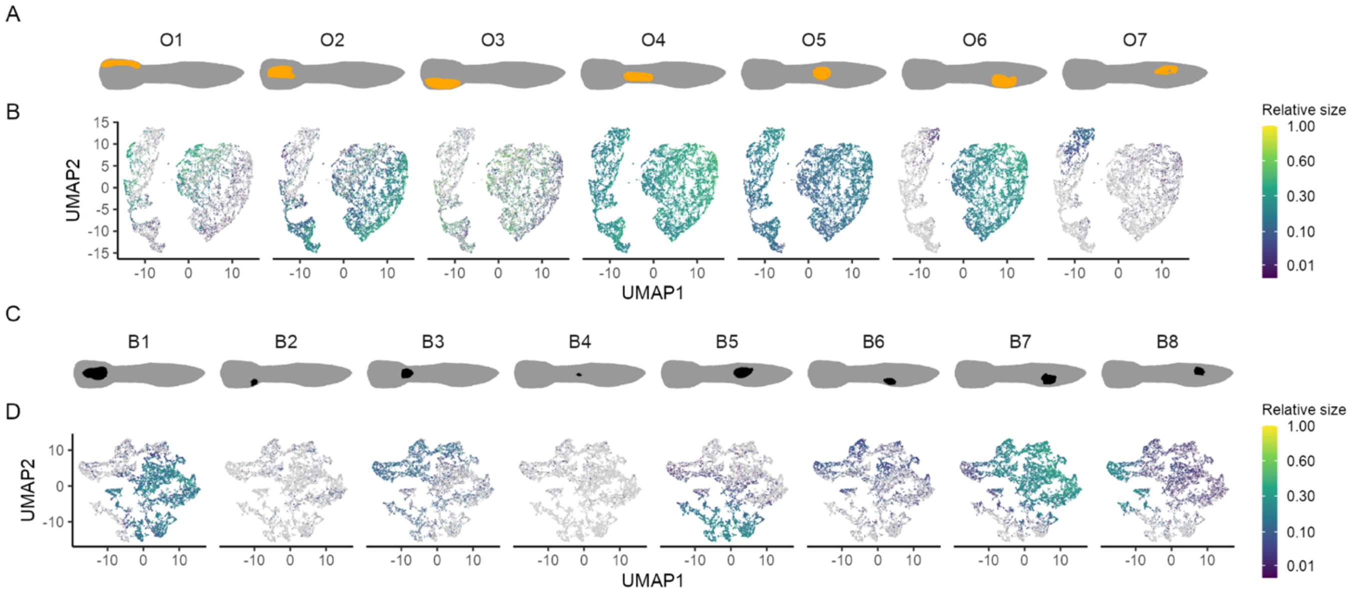


**Figure S17: Pattern space encodes variation in the incidence and size of ornaments.** A&C) Pictograms of seven orange and eight black ornaments, matching with the associated panels in B&D respectively. B&D) Axes are the UMAP reduced representation of five-dimensional pattern space. Individuals are colored by the size of their relevant ornament, expressed as a fraction of the largest observed size, or colored grey if they lack the ornament. Note the use of a logarithmic scale for ornament size.

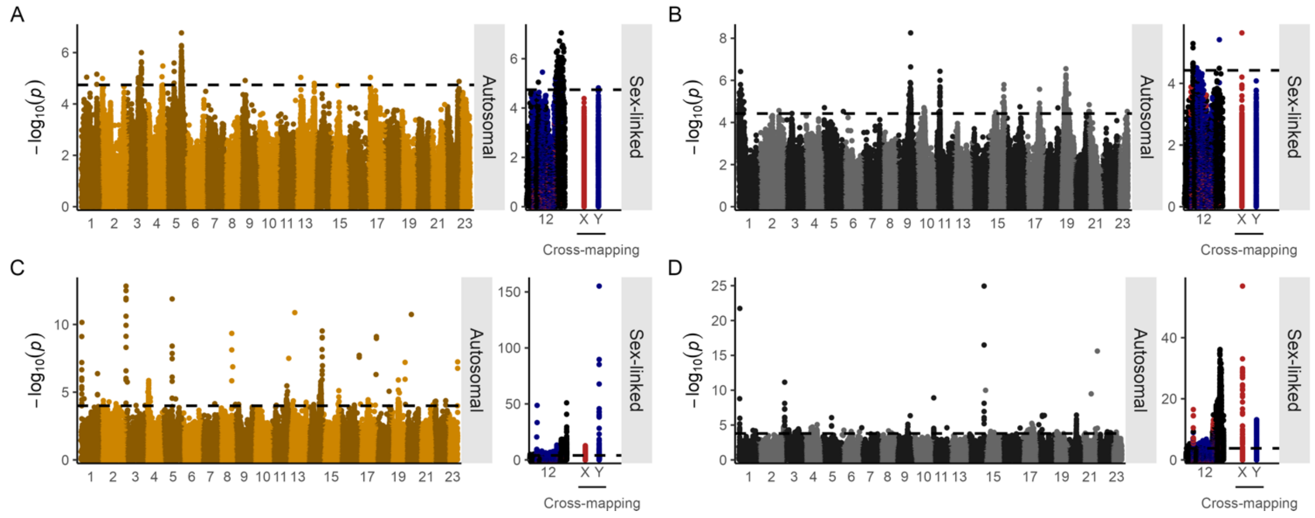


**Fig S18: Manhattan plots of genomic associations with orange and black color, mapped to a male reference (*37*).** A) Results of multivariate GWAS on position in orange (left) and black (right) patternspace. Points represent SNPs and small indels, with their genomic location on the x-axis, and the p-value of the association on the y-axis. Numbers along the x axis denote linkage groups, with unplaced scaffolds plotted on the right-hand side. Significant associations (5% FDR) are colored by their inferred pattern of inheritance, whereas non-significant associations are colored in alternating shades of grey. B) GWAS results for the association with the presence/absence of select ornaments: O5, O1, and O6 (left), and B1, B5 and B8 (right).

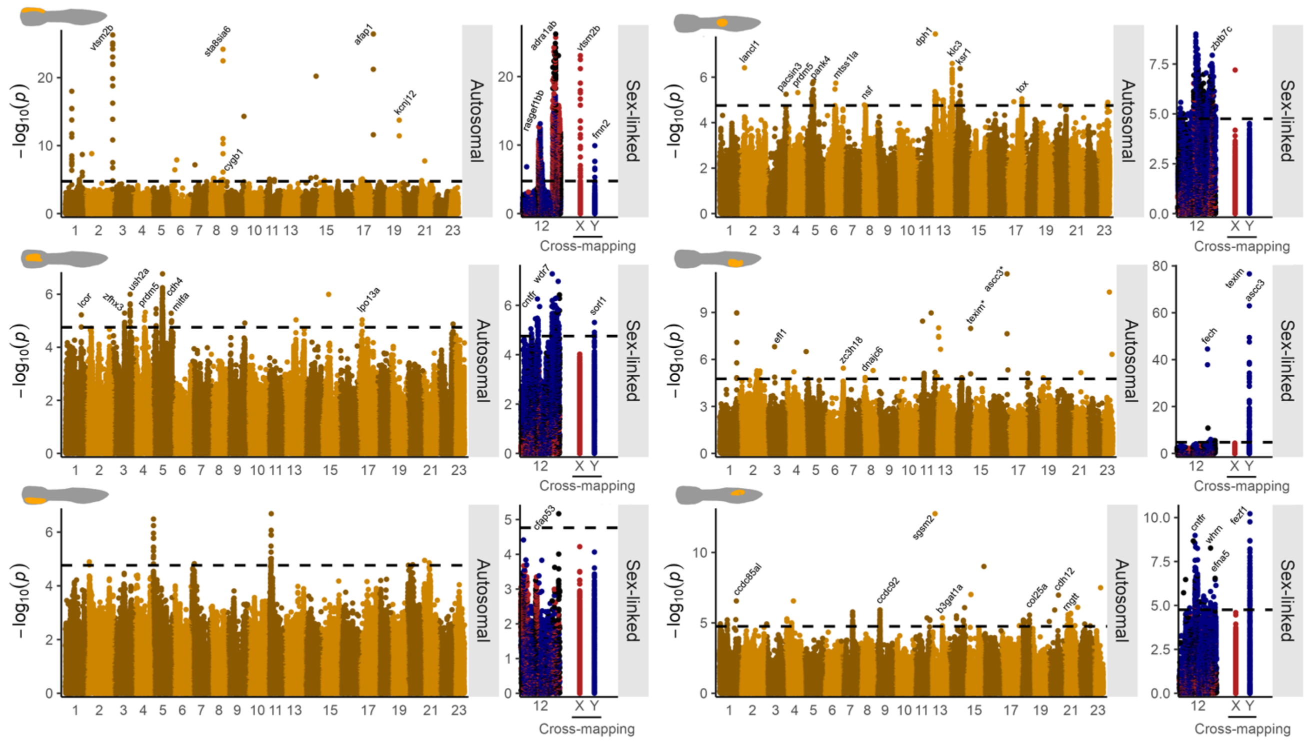

**Fig S19: GWAS results for the presence/absence of six orange ornaments**, as in Fig 5. Points represent SNPs and small indels, with their genomic location on the x-axis, and the p-value of the association on the y-axis. Numbers along the x axis denote linkage groups. Sex-linked loci are placed to the side, with both the sex-chromosome (LG12) and sex-linked loci cross-mapping to the autosomes shown, colored by their putative origin (red = X, blue = Y). Peaks are labelled with gene names, but unnamed and uncharacterized genes are shown. Names with asterisk (*) indicate that more significant cross-mapping SNPs were present in the same peak..

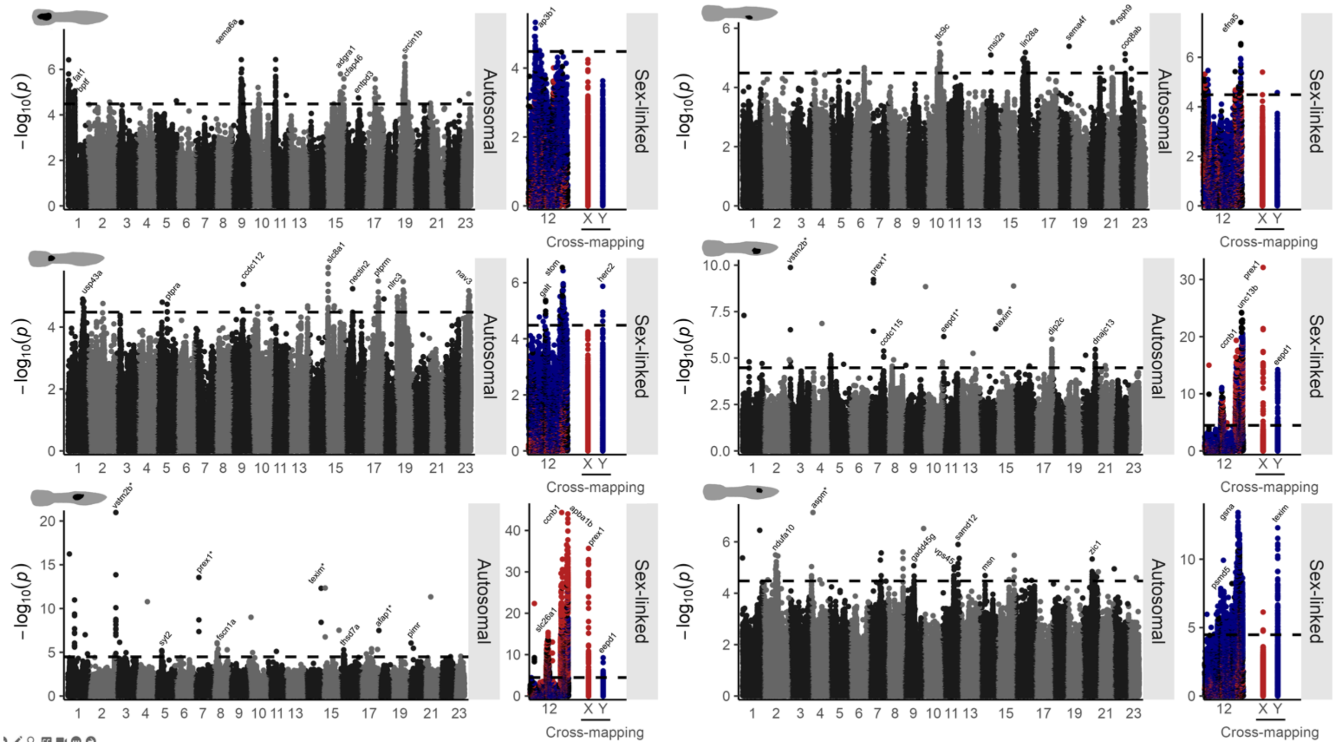

**Fig S20: GWAS results for the presence/absence of six black ornaments**, as in Fig 5. Points represent SNPs and small indels, with their genomic location on the x-axis, and the p-value of the association on the y-axis. Numbers along the x axis denote linkage groups. Sex-linked loci are placed to the side, with both the sex-chromosome (LG12) and sex-linked loci cross-mapping to the autosomes shown, colored by their putative origin (red = X, blue = Y). Peaks are labelled with gene names, but unnamed and uncharacterized genes are shown. Names with asterisk (*) indicate that more significant cross-mapping SNPs were present in the same peak.

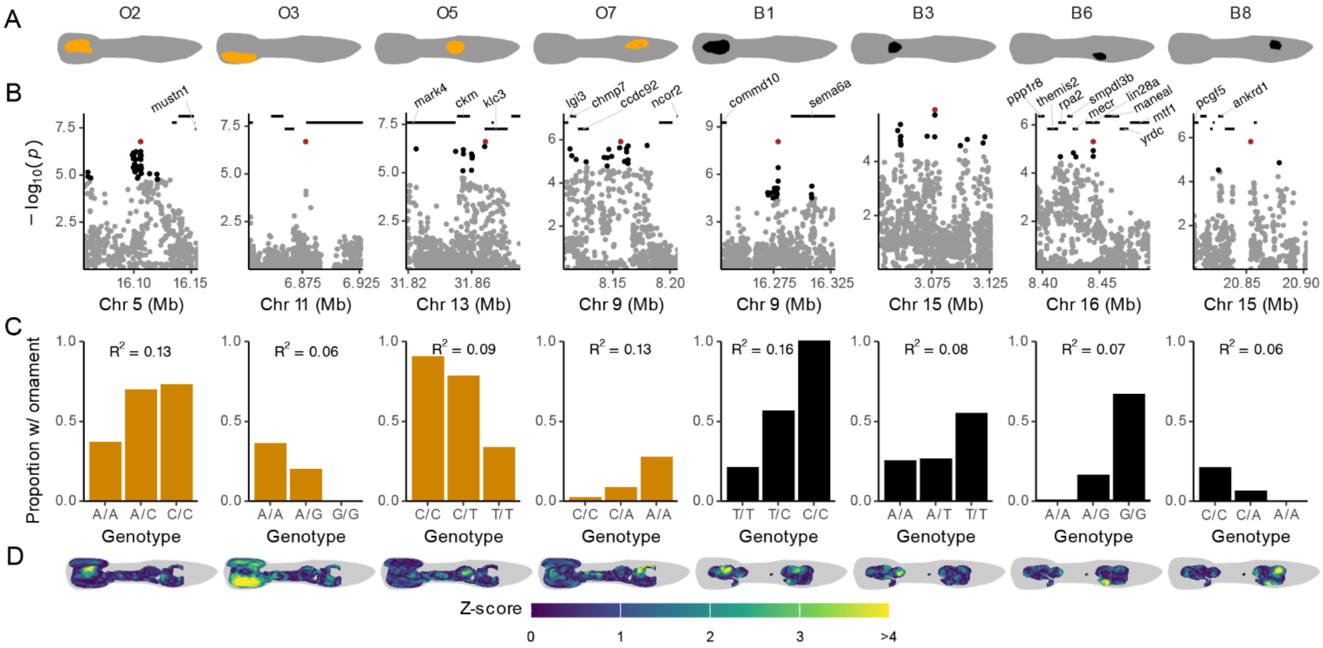


**Figure S21: Autosomally inherited ornaments are complex traits.** A) Pictograms of eight ornaments with (partial) autosomal inheritance. B) Illustrative GWAS peaks for the association with the ornaments depicted in A. Each point is a genetic variant, with black points surpassing the FDR-threshold, and the red point is the top variant. Horizontal lines show the location of all annotated genes, with labels only shown for characterized genes. C) Bars indicate the proportion of individuals with the ornament (depicted in A) depending on their genotype at the top variant. The proportion of variation in the phenotype that is explained by the top variant is displayed as Nagelkerke's $R^2$ calculated from a logistic regression. D) PheWAS heatmaps displaying Z-scores for the effect of the top variant on orange or black color across the body, controlling for autosomal and sex-linked relatedness.

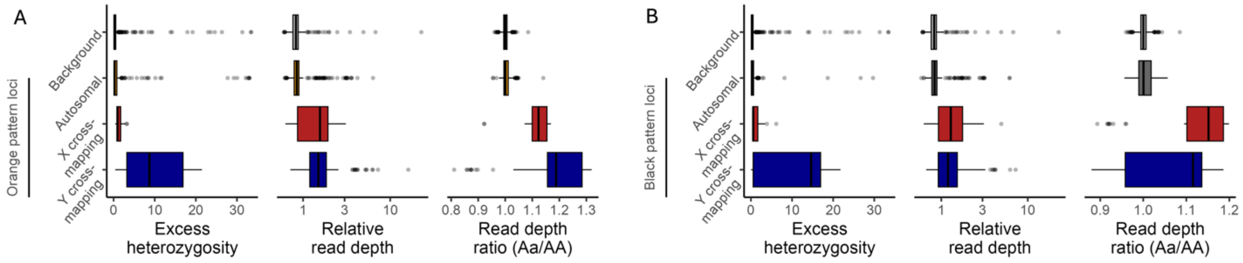


**Figure S22: Loci with sex-linked patterns of inheritance mapping to autosomes show evidence of cross-mapping.** A) Excess heterozygosity (-$\log_{10}$ $p$-value of $\chi^2$ test against HWE prediction), read depth (relative to genomic average), and read depth ratio (heterozygous read depth over homozygous reference read depth) are plotted for four groups of loci. Background loci are 1,000 randomly sampled SNPs from the autosomes. Orange pattern loci are all autosomally mapping, but separated

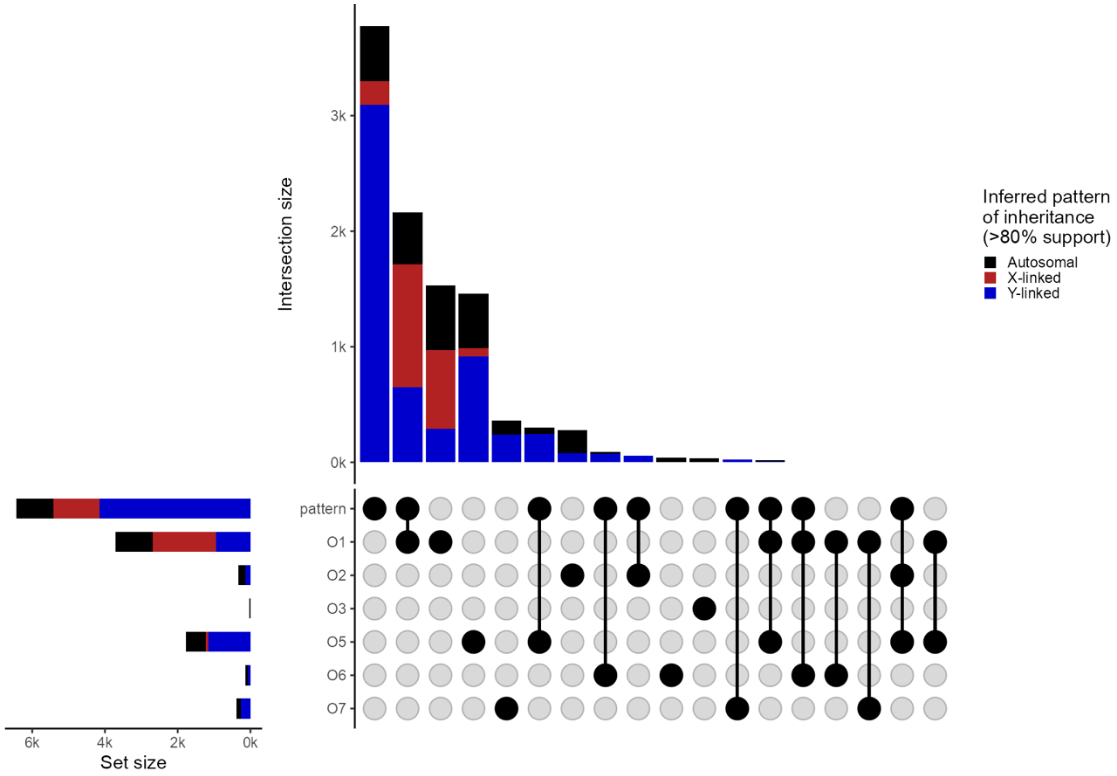


**Fig S23: Orange pattern elements have largely independent genetic architectures.** Upset plot showing the unique and overlapping elements in significant variants detected in the GWAS for orange pattern and six orange ornaments, across the whole genome. Vertical bars represent the count of elements unique or shared between sets, with the connected dot display indicating the specific sets involved in each intersection. Horizontal bars show individual set sizes. Bars are colored proportionally by the inferred inheritance pattern of the variants in that set or intersection.

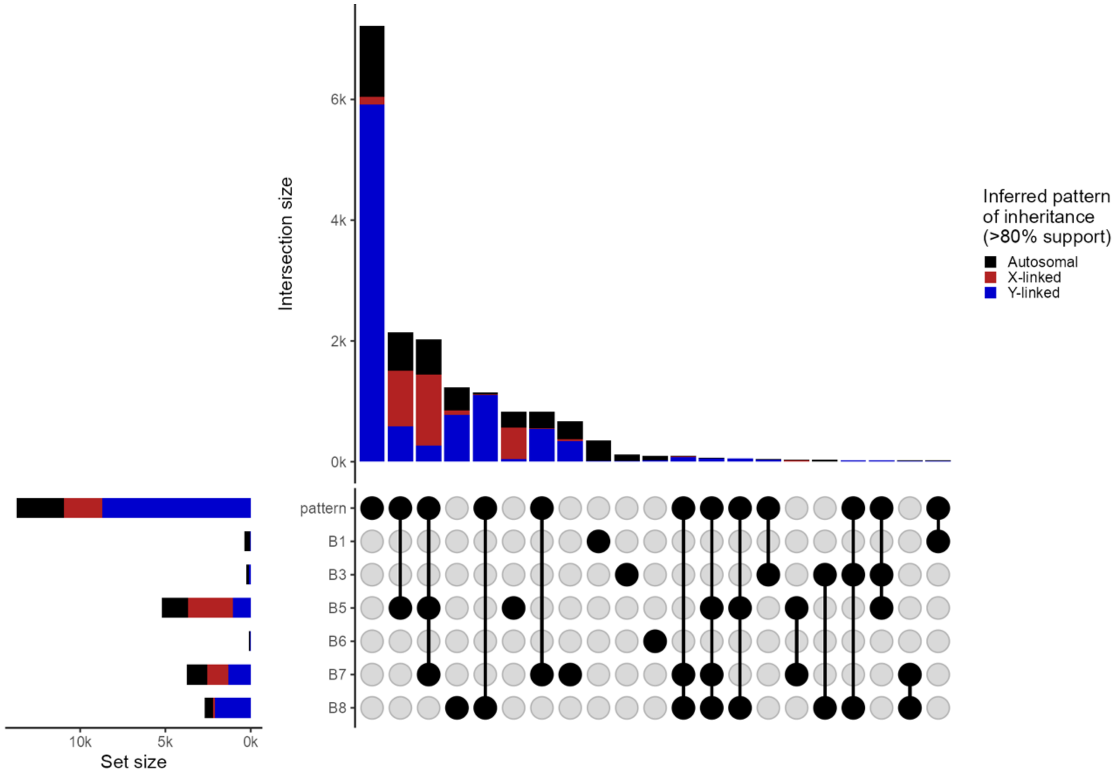


**Fig S24: Black pattern elements have largely independent genetic architectures.** Upset plot showing the unique and overlapping elements in significant variants detected in the GWAS for black pattern and six black ornaments, across the whole genome. Vertical bars represent the count of elements unique or shared between sets, with the connected dot display indicating the specific sets involved in each intersection. Horizontal bars show individual set sizes. Bars are colored proportionally by the inferred inheritance pattern of the variants in that set or intersection.

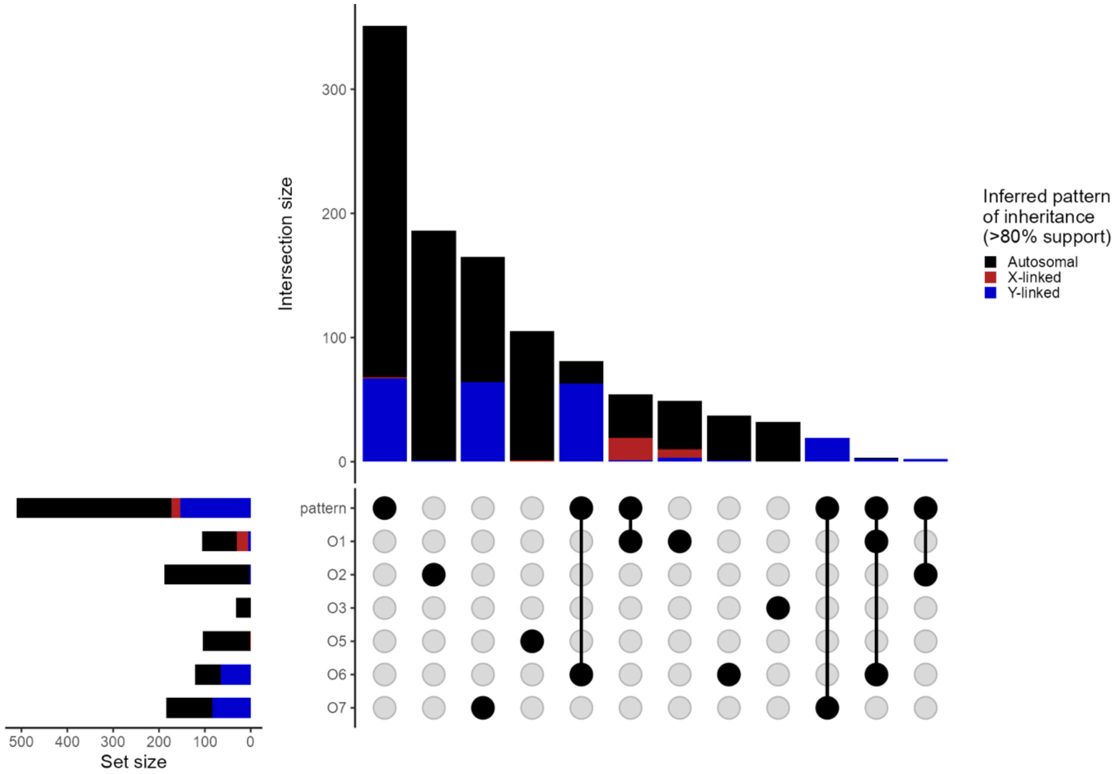


**Fig S25:** Upset plot showing the unique and overlapping elements in significant variants detected in the GWAS for orange pattern and six orange ornaments, for variants mapping to the autosomes. Vertical bars represent the count of elements unique or shared between sets, with the connected dot display indicating the specific sets involved in each intersection. Horizontal bars show individual set sizes. Bars are colored proportionally by the inferred inheritance pattern of the variants in that set or intersection.

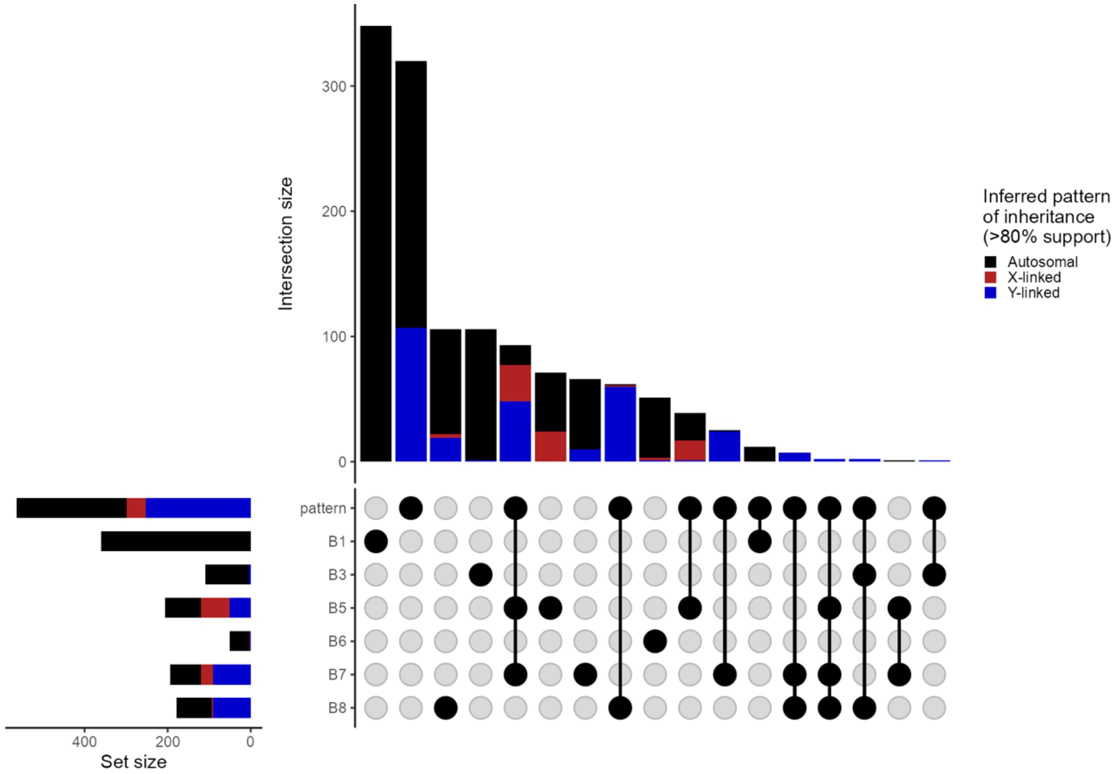


**Fig S26:** Upset plot showing the unique and overlapping elements in significant variants detected in the GWAS for black pattern and six black ornaments, for variants mapping to the autosomes. Vertical bars represent the count of elements unique or shared between sets, with the connected dot display indicating the specific sets involved in each intersection. Horizontal bars show individual set sizes. Bars are colored proportionally by the inferred inheritance pattern of the variants in that set or intersection.

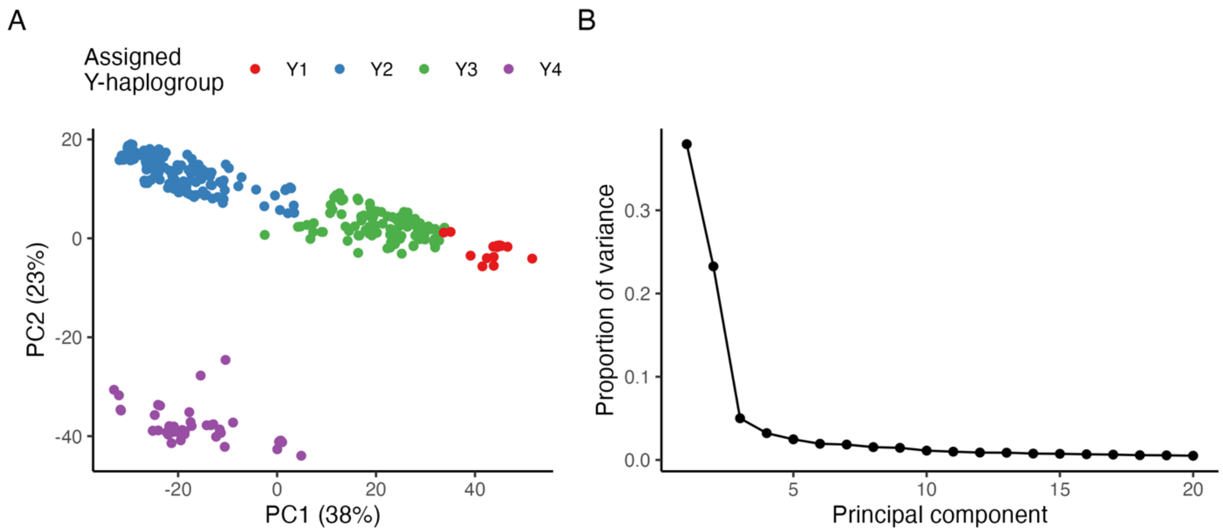


**Fig S27:** Principal component analysis of variants associated with orange pattern and Y-linked inheritance pattern. A) shows the biplot with labeled inferred Y-haplogroups, while B) shows the scree plot.

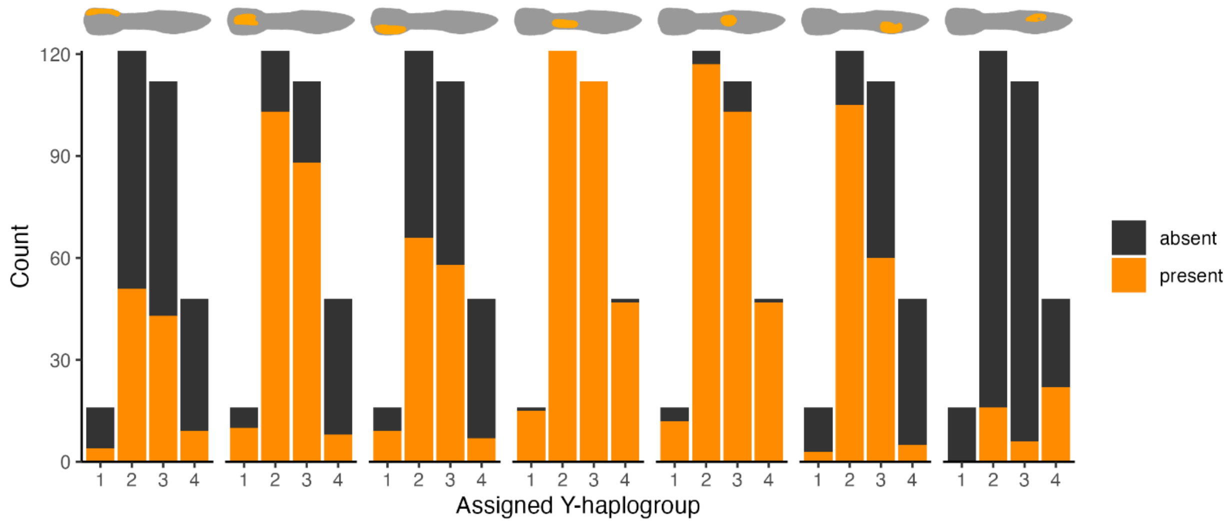


**Fig S28**: Stacked bar chart showing the relationship between Y-haplogroup and ornament presence.

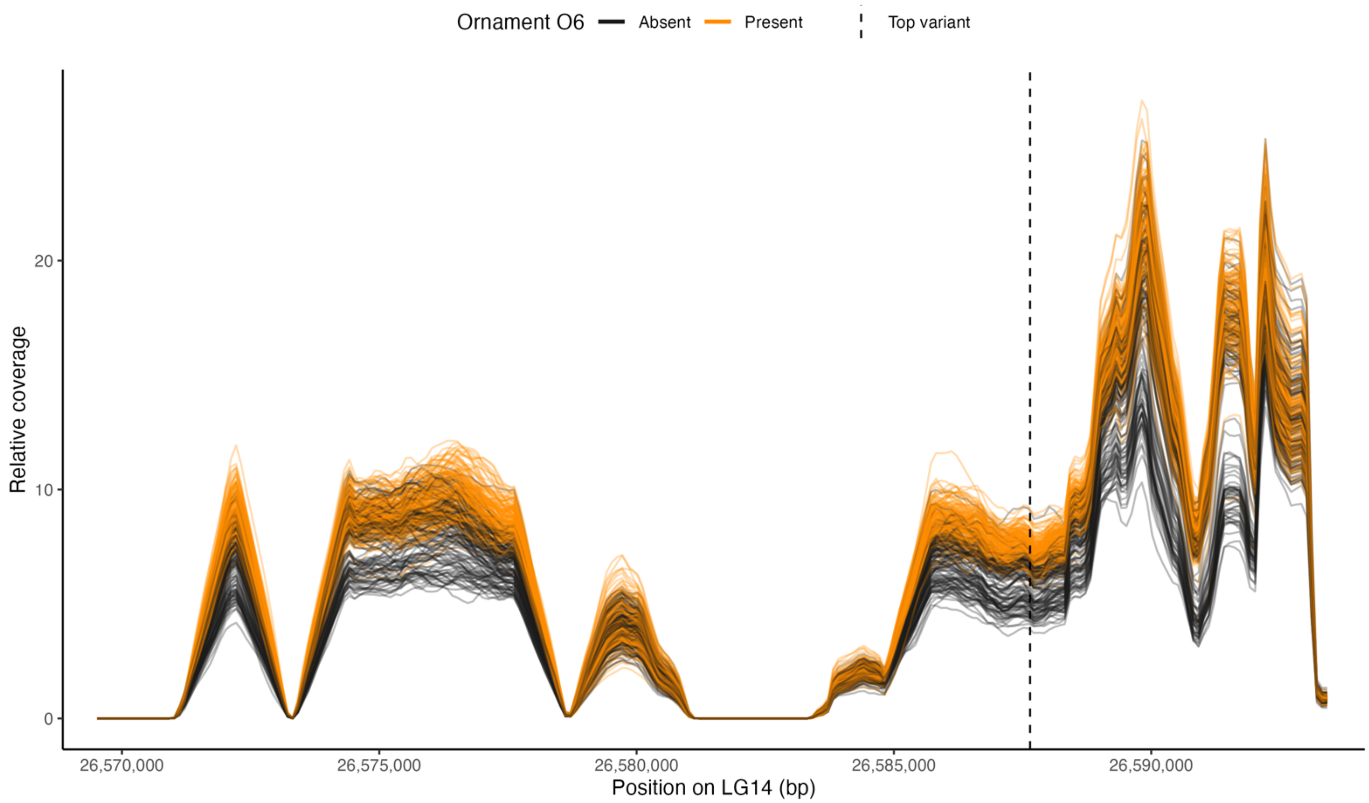


**Fig S29: Coverage in the region near *texim* on LG14.** Coverage is plotted relative to the genome wide sample average (= 1), calculated in moving windows of 1000bp at a distance 100bp. Each line is 1 male sample, colored by whether the male possesses ornament O6. The location of the top variant from the genomic association with orange color pattern is indicated with the dotted vertical line.

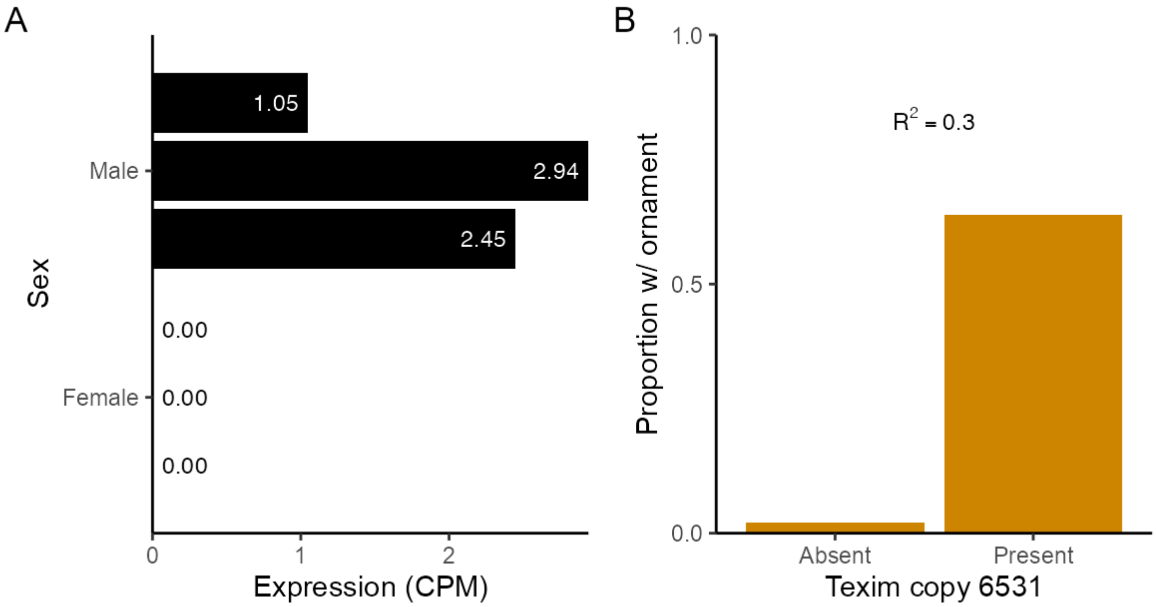


**Figure S30: Expression near *texim* and relation to ornament O6.** A) Expression of *texim* copy 6531 in three male and three female pools, as counts per million (CPM). B) The proportion of males with ornament O6, as a function of the presence of *texim* copy 6531, as determined by reads with diagnostic SNPs for that copy. The proportion of phenotypic variance is sh down as Nagelkerke's $R^2$.

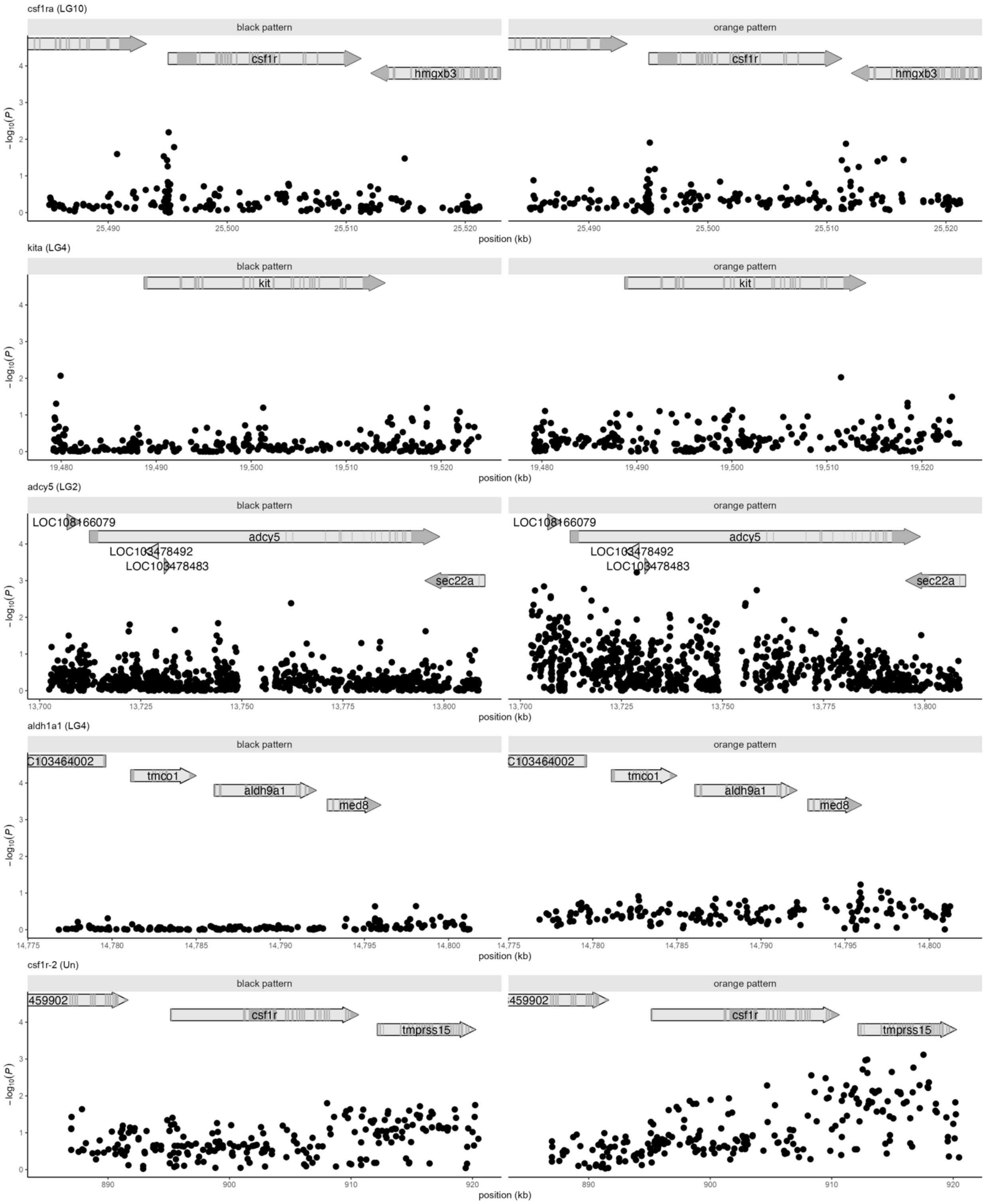


**Figure S31**: No relationship between black and orange color pattern and genes previously indicated in carotenoid and melanic pigmentation. Panels show GWAS results for black and orange pattern space.

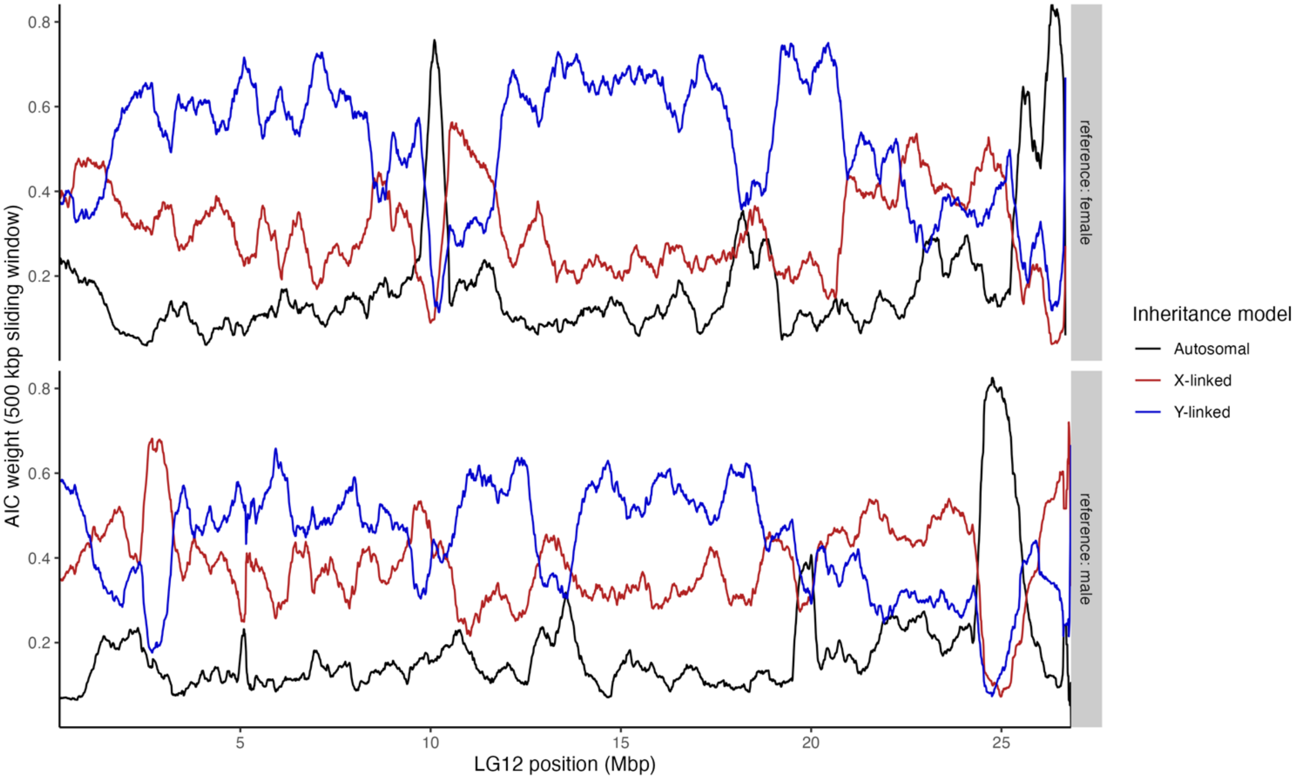


**Figure S32: Inferred patterns of inheritance for 100,000 variants along LG12**, for both a female and male reference genome. The AIC weight (relative likelihood) for each inheritance model is averaged across sliding windows of 500 kbp. Note that for the female reference, the autosomally inherited portion around 10 Mbp is a likely assembly error, as this region is placed on LG7 in the male reference. Note that for the male reference, the LG12 assembly includes both putative X and Y scaffolds.

Supplemental tables

Table S1: Architecture of the convolutional neural net used in step 3 of the pipeline.

| Layer | Activation | Filters | Output shape | Spatial dropout |
|---|---|---|---|---|
| Convolution | Relu | 32 | [400, 144, 32] | 0.4 |
| Convolution | Relu | 64 | [400, 144, 64] | 0.4 |
| Max pooling | | | [200, 72, 64] | |
| Convolution | Relu | 128 | [200, 72, 128] | 0.4 |
| Deconvolution | Relu | 128 | [200, 72, 128] | 0.4 |
| Upsampling | | | [400, 144, 128] | |
| Deconvolution | Relu | 64 | [400, 144, 64] | 0.4 |
| Deconvolution | Linear | 4 | [400, 144, 4] | |
